# NFATc2 enhances tumor-initiating phenotypes through the NFATc2/SOX2/ALDH axis in lung adenocarcinoma

**DOI:** 10.1101/131987

**Authors:** Zhi-Jie Xiao, Jing Liu, Si-Qi Wang, Yun Zhu, Xu-Yuan Gao, Vicky Pui-Chi Tin, Jing Qin, Jun-Wen Wang, Maria Pik Wong

## Abstract

Cancers display intratumoral genetic and molecular heterogeneity with tumor initiating cells (TIC) showing enhanced tumor phenotypes. In this study, we show the calcium pathway transcription factor NFATc2 is a novel regulator of lung TIC through the NFATc2/SOX2/ALDH1A1 regulatory axis. *In vitro* and *in vivo* cancer cell modeling demonstrated supportive evidences including cell renewal, tumorigenicity at limiting dose, cell motility, resistance to cytotoxic chemotherapy and EGFR targeted therapy. In human lung cancers, high NFATc2 expression predicts poor tumor differentiation, adverse recurrence-free and overall patient survivals. Mechanistic investigations identified NFATc2 response elements in the SOX2 3’ enhancer region, and NFATc2/SOX2 coupling upregulates ALDH1A1 by binding to its 5’ enhancer. Through this axis, oxidative stresses and reactive oxygen species induced by cancer drug treatment are attenuated, accounting for a mutation-independent mechanism of drug resistance. Targeting this axis provides a novel approach for the long term treatment of lung cancer through TIC elimination.

## Introduction

Lung cancer results from mutations induced by DNA adducts, free radicals and reactive oxygen species (ROS) generated during tobacco smoking and chronic inflammation (Acharya, Das et al. 2010, Okumura, Yoshida et al. 2012, Houghton 2013). Late presentation and the lack of effective long term therapy accounts for the high mortality necessitating new therapy. Recent research has shown the cellular landscape of a cancer is heterogeneous. Cells showing aberrant expressions of various molecules have enhanced propensities for survival, tumorigenicity, drug resistance and are designated as cancer stem cells or tumor-initiating cells (TIC). In some cancers, constitutive activities of inherent embryonic or developmental pathways for stem cell renewal are involved in TIC maintenance, such as the WNT/β-catenin pathway in colonic adenocarcinomas (AD). In tissues with slow cell turnover, mechanisms that elicit cell plasticity and stemness properties during tissue response to intracellular and extracellular stresses are involved (Valent, Bonnet et al. 2012, Visvader and Lindeman 2012, Beck and Blanpain 2013). For the adult lung, stem cell niches or their physiological regulatory mechanisms are ill-defined, and molecular programs sustaining lung TIC are still elusive.

Intracellular free calcium is at the hub of multiple interacting pathways activated by extracellular and/or intrinsic stimulations, e.g. EGFR, endoplasmic reticulum and mitochondrial stresses, etc. (Roderick and Cook 2008, Prevarskaya, Skryma et al. 2010, Zhao, Wang et al. 2013, Deliot and Constantin 2015), raising the possibility stress signals transduced by calcium pathway mediators could be involved in the induction of TIC phenotypes. In cancers of the breast, pancreas, colon and melanoma, the calcium signaling transcription factor Nuclear Factor of Activated T-cells (NFAT) has been shown to contribute to malignant properties including cell invasion, migration, survival, proliferation, stromal modulation and angiogenesis (Werneck, Vieira-de-Abreu et al. 2011, Gerlach, Daniel et al. 2012, Qin, Nag et al. 2014). But information on its role in TIC phenotypes especially drug resistance is limited. Details of the molecular pathways linking calcium signaling to TIC induction and drug resistance have not been reported. In this study, we demonstrated the isoform NFATc2 supports tumorigenicity, cell survival, motility and drug resistance of human lung AD. Amongst essential factors of pluripotency, SOX2 is an NFATc2 target upregulated through its 3’ enhancer, while SOX2 couples to a 5’ enhancer of ALDH1A1 mediating overexpression. NFATc2 induces TIC that coexpress the markers ALDH and CD44 (ALDH^+^/CD44^+^-TIC), where ROS scavenging by the NFATc2/SOX2/ALDH1A1 axis enhances drug resistance. Our study reports a novel mechanism of lung TIC maintenance pathway whereby micro-environmental stimulation is linked to induction of stemness phenotypes, evasion of cell death and enhancement of drug resistance in lung AD. NFATc2 could be an important target in treatment strategies aiming at disruption of TIC in lung cancer.

### Results

### NFATc2 expression correlated with adverse survivals of human NSCLC

NFATc2 transcripts were significantly overexpressed in NSCLC compared to normal lung (Fig. 1A). In 102 excised primary human lung cancers expressing NFATc2 by IHC, 41 (40.2%) showed high level activated NFATc2 shown as intense nuclear staining in large sheets of tumor cells distributed over extensive areas, while 61 (59.8%) showed low expression featuring weak cytoplasmic with or without nuclear staining in scattered, isolated or small clusters of tumor cells (Fig. 1B, C). In normal lung epithelium, NFATc2 was expressed in the bronchiolar stem cell compartment of basal reserve cells while differentiated bronchiolar cells or alveolar pneumocytes were negative (Fig. 1D). Using log rank test and Kaplan-Meier survival analysis, tumors with high level NFATc2 expression showed significantly shorter recurrence-free survival (RFS) and overall survival (OS) (Fig. 1C, D). High NFATc2 expression significantly predicted poor tumor differentiation, advanced tumor stage and TNM stage (Table. 1A). Furthermore, multivariate Cox regression analysis showed high NFATc2, late pathological stage, age and smoking history were independent prognostic indicators for shorter OS, while high NFATc2 and advanced pathological stage were associated with shorter RFS (Table. 1B, C). The results indicated NFATc2 expression is associated with repressed tumor differentiation and adverse patient outcomes.

**Figure 1.**
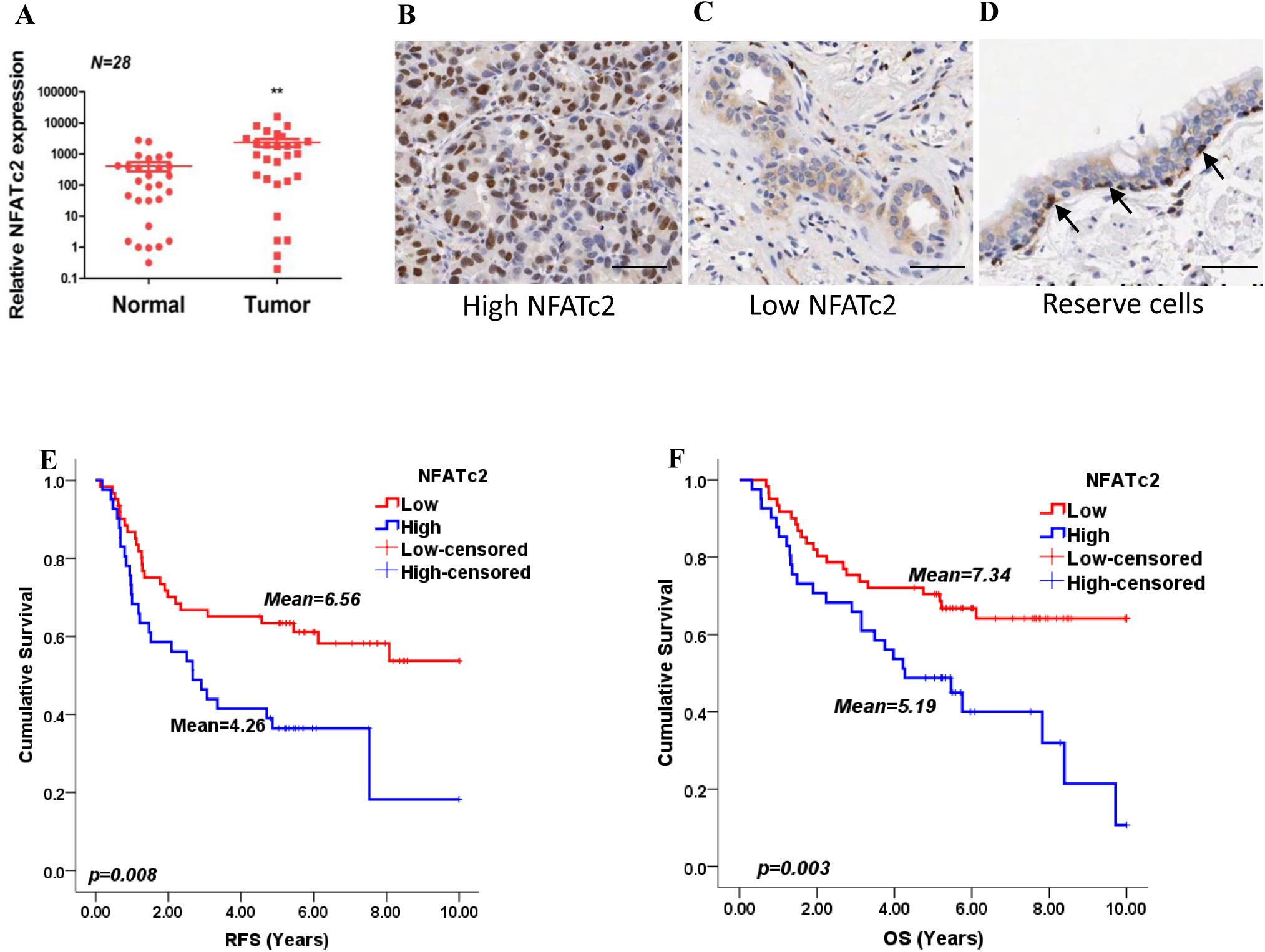
NFATc2 overexpressed in human NSCLC. **(A)** NFATc2 expression analyzed by qPCR human NSCLC and corresponding normal lung. P: Wilcoxon test. P= 0.0003 (**B-C)** NFATc2 expression in lung cancers was analyzed by IHC, with representative views of high NFATc2 score featuring strong nuclear staining in the majority of cancer cells shown in (B), and low NFATc2 score featuring weak nuclear staining in scattered cancer cells shown in (C), respectively. Scale bars, 50 µM. **(D)** NFATc2 expression in normal bronchial epithelium by IHC. Nuclear NFATc2 was expressed in the reserve cell layer of normal bronchial epithelium. Scale bars, 50 µM. (**E-F)** Log-rank tests and Kaplan Meier survival curves for recurrence-free survival (RFS) (E) and overall survival (OS) (F) on 102 resected primary NSCLC stratified by NFATc2 expression score. Figure1-source data 1: Statistical analyses for figure 1A.

**Table 1.**
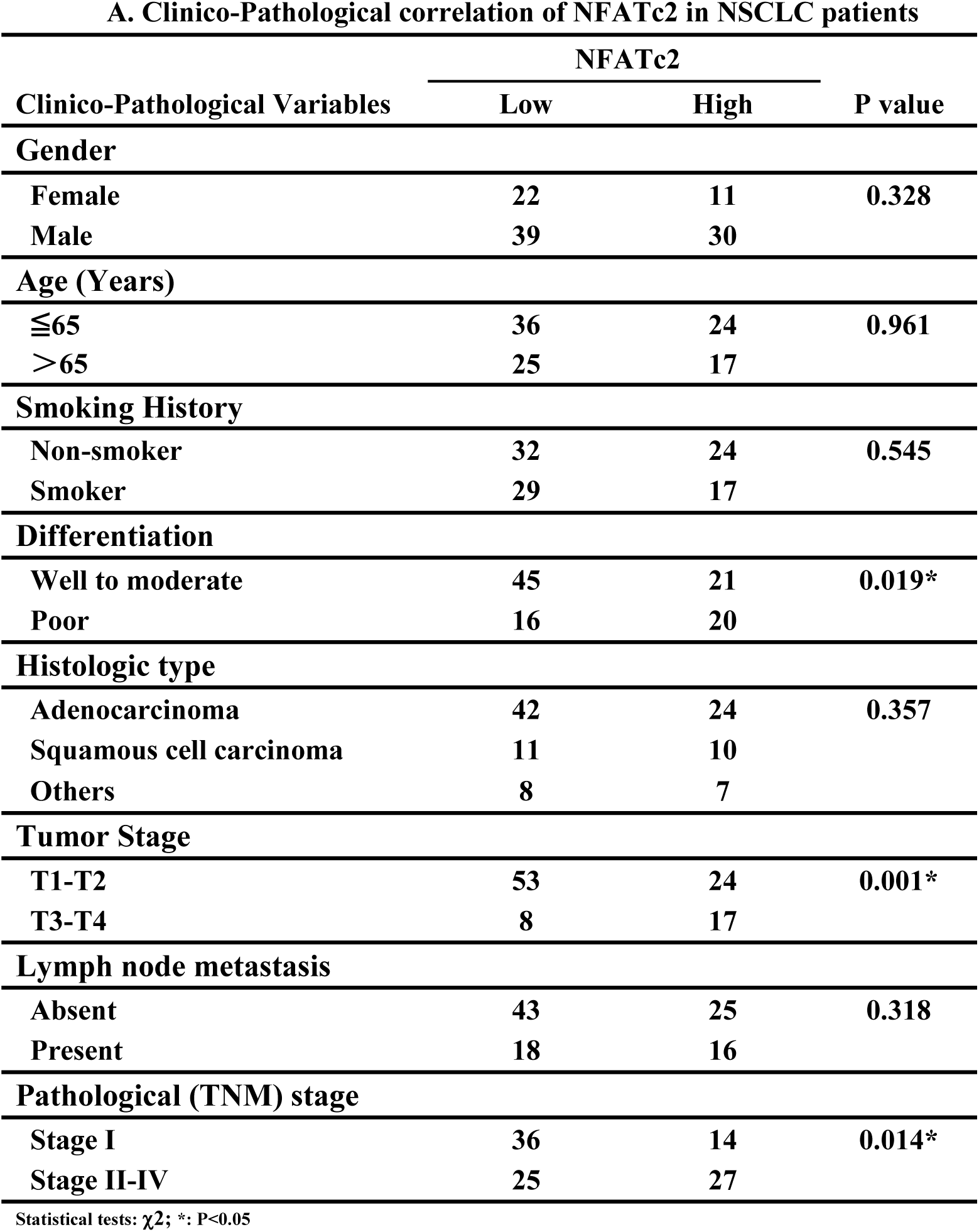
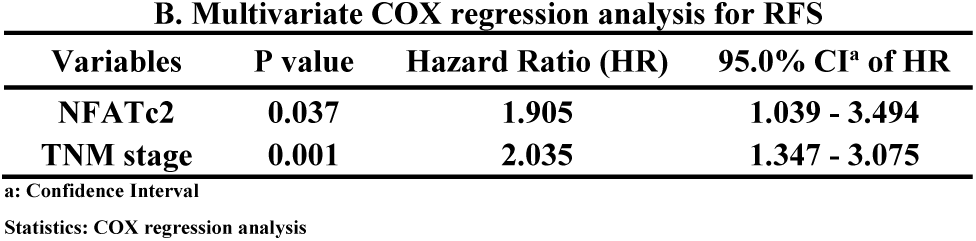
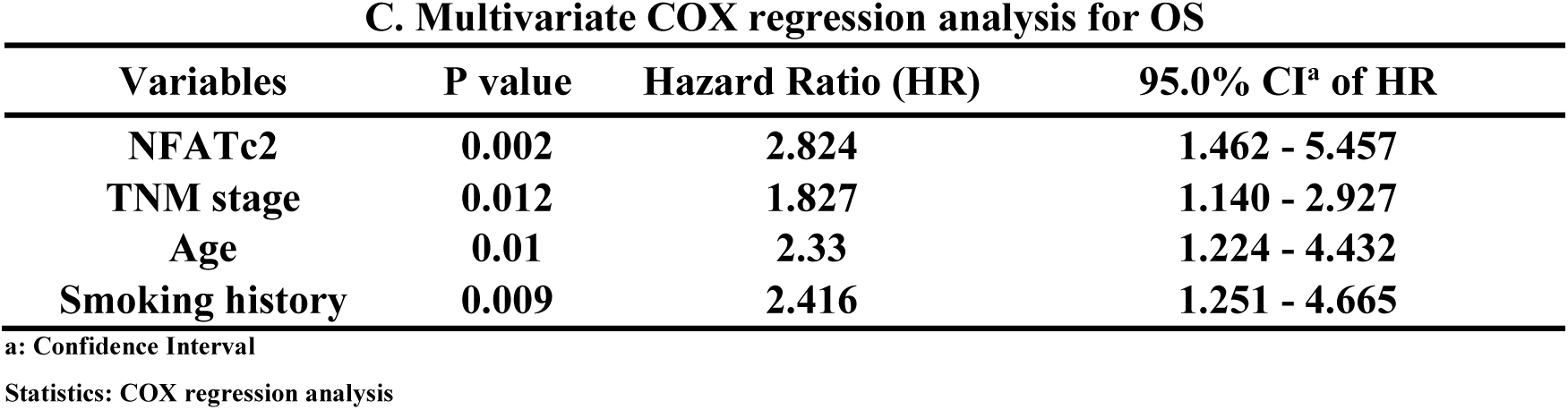
A. Clinico-Pathological correlation of NFATc2 in NSCLC patients

| Clinico-Pathological Variables | NFATc2 |  | P value |
| --- | --- | --- | --- |
|  | Low | High |  |
| Gender |  |  |  |
| Female | 22 | 11 | 0.328 |
| Male | 39 | 30 |  |
| Age (Years) |  |  |  |
| ≤65 | 36 | 24 | 0.961 |
| >65 | 25 | 17 |  |
| Smoking History |  |  |  |
| Non-smoker | 32 | 24 | 0.545 |
| Smoker | 29 | 17 |  |
| Differentiation |  |  |  |
| Well to moderate | 45 | 21 | 0.019* |
| Poor | 16 | 20 |  |
| Histologic type |  |  |  |
| Adenocarcinoma | 42 | 24 | 0.357 |
| Squamous cell carcinoma | 11 | 10 |  |
| Others | 8 | 7 |  |
| Tumor Stage |  |  |  |
| T1-T2 | 53 | 24 | 0.001* |
| T3-T4 | 8 | 17 |  |
| Lymph node metastasis |  |  |  |
| Absent | 43 | 25 | 0.318 |
| Present | 18 | 16 |  |
| Pathological (TNM) stage |  |  |  |
| Stage I | 36 | 14 | 0.014* |
| Stage II-IV | 25 | 27 |  |

Statistical tests: $\chi^2$ ; \*: P<0.05**B. Multivariate COX regression analysis for RFS**
| Variables | P value | Hazard Ratio (HR) | 95.0% CI<sup>a</sup> of HR |
| --- | --- | --- | --- |
| NFATc2 | 0.037 | 1.905 | 1.039 - 3.494 |
| TNM stage | 0.001 | 2.035 | 1.347 - 3.075 |

<sup>a</sup>: Confidence Interval

C. Multivariate COX regression analysis for OS
| Variables | P value | Hazard Ratio (HR) | 95.0% CI<sup>a</sup> of HR |
| --- | --- | --- | --- |
| NFATc2 | 0.002 | 2.824 | 1.462 - 5.457 |
| TNM stage | 0.012 | 1.827 | 1.140 - 2.927 |
| Age | 0.01 | 2.33 | 1.224 - 4.432 |
| Smoking history | 0.009 | 2.416 | 1.251 - 4.665 |

a: Confidence Interval
Statistics: COX regression analysis

### NFATc2 enhanced *in vitro* and *in vivo* tumorigenesis and cell motility

If NFATc2 supports tumor initiating phenotypes, it is expected to be expressed at a higher level in TIC compared to non-TIC. Marker-independent TIC surrogates were first raised as tumorspheres from 4 lung AD cell lines cultured in non-adherent, stem cell conditions (Liu, Deng et al. 2007, Shi, Li et al. 2015, Sun, Hu et al. 2015). Compared to non-TIC harvested from the monolayers, tumorspheres expressed higher NFATc2 by Western blot (Fig. 2A), while transcripts of *NFATc2* and its target *FASL* were also significantly upregulated (Figure 2-figure supplement 1A). Furthermore, luciferase reporter assays showed significantly higher NFAT activities in spheres isolated from H1299 and A549 cells (Figure 2-figure supplement 1B).

**Figure 2.**
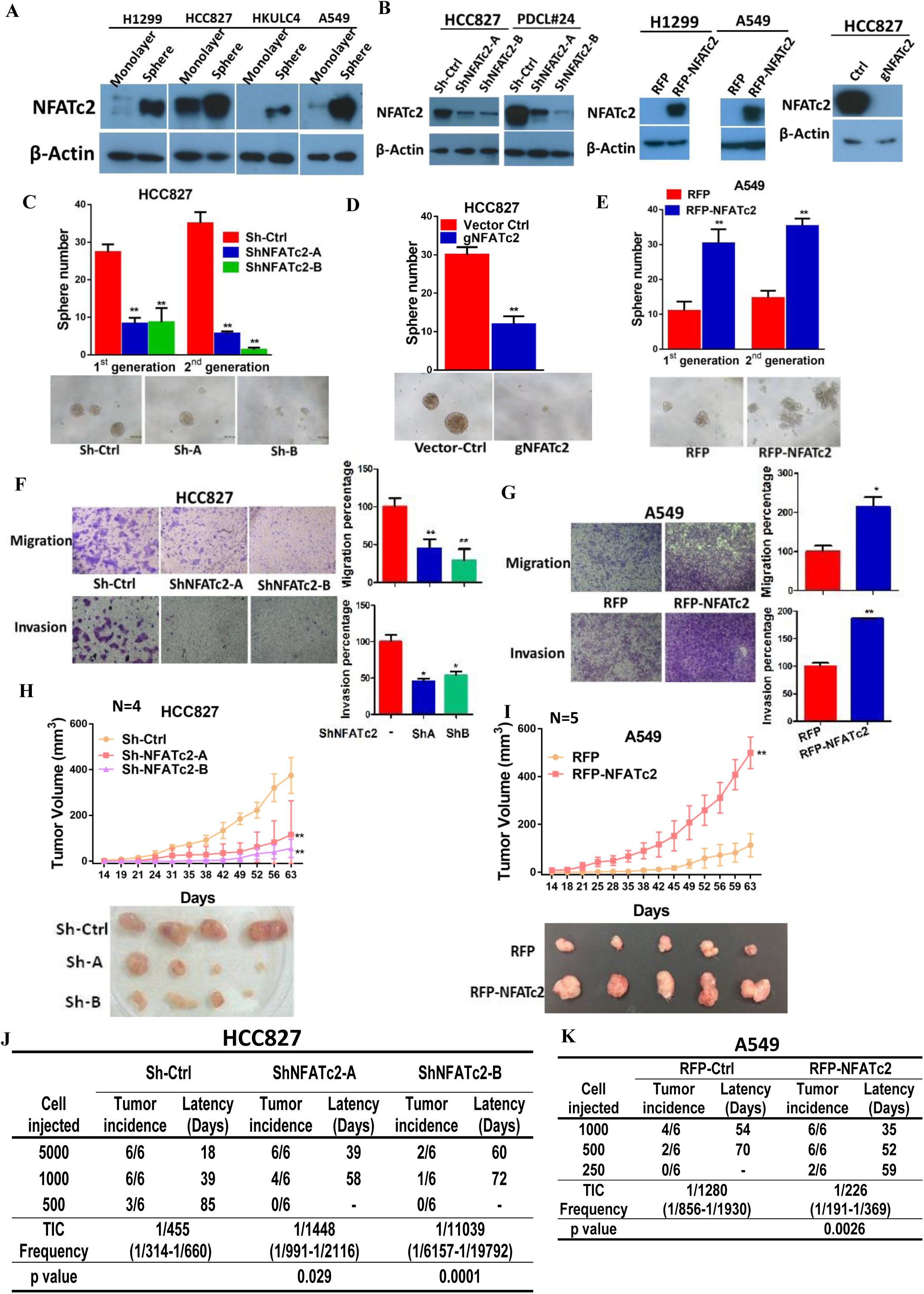
NFATc2 regulated lung TIC phenotypes. **(A)** Expression of NFATc2 analyzed by Western blot in TIC isolated by tumorspheres compared to the corresponding monolayer controls. (**B)** Expression of NFATc2 analyzed by Western blot in cells with stable NFATc2 knockdown, overexpression, or knockout, respectively. (**C-E)** Tumorsphere formation and serial passage assays in HCC827 cells after stable NFATc2 knockdown (C) or knockout (D), or in A549 cells with stable NFATc2 over-expression (E) compared to controls. (**F-G)** Cell migration and invasion assays in cells with stable NFATc2 knock down (F) or over-expression (G). C-G: *p<0.05 **p<0.01, comparison with control by t-test. Error bar indicates the mean ± SD for at least three independent replicates. (**H-I)** 1×10^4^ of HCC827 cells (H) and A549 cells (I) were subcutaneously inoculated into the flanks of SCID mice, and tumor volumes were monitored. Representative tumor images and tumor growth curves are shown. **p<0.0001, comparison with respective control by two-way ANOVA. Error bar indicates the mean ± SD of tumor volumes of mice as indicated. (**J-K)** Limiting dilution assay *in vivo*. Indicated numbers of HCC827cells (J) and A549 cells (K) were subcutaneously inoculated into SCID mice, and the tumor incidence and latency were monitored for 3 months. The TIC frequency and P values were calculated using the L-Calc software (Stemcell Tech, Vancouver, Canada, http://www.stemcell.com). The following figure supplements and source data are available for figure 2: Figure 2-figure supplement 1: NFATc2 was up-regulated in tumorspheres. Figure 2-figure supplement 2: NFATc2 regulated *in vitro* TIC properties. Figure 2-figure supplement 3: NFATc2 regulated *in vivo* tumorigenesis. Figure 2-source data 1: Statistical analyses for figure 2H and I, and figure supplement 3A.

Next, two lung cancer cell lines with high basal NFATc2 (HCC827, PDCL#24) were silenced by 2 shRNA sequences (shNFATc2-A and -B), and two cell lines with relatively low *de novo* expression (A549, H1299) were used for NFATc2 ectopic expression (Fig. 2B). NFATc2 knockout by the CRISPR/CAS9 technique using gRNA targeting NFATc2 (gNFATc2) was also performed on HCC827 cells (Fig. 2B). Abrogation of NFATc2 significantly reduced 60-70% of tumorspheres in all the cell models and inhibited tumorspheres renewability for 2 consecutive generations (Fig. 2C-D, figure 2-figure supplement 2A), while overexpression significantly augmented tumorspheres in both A549 and H1299 (Fig. 2E, figure 2-figure supplement 2B). Transwell assays for cell motility showed silencing NFATc2 significantly reduced the migration and invasion ability of both HCC827 and PDCL#24 cells (Fig. 2F, figure 2-figure supplement 2C), while the opposite effects were rendered by NFATc2 overexpression in A549 and H1299 cells (Fig. 2G, figure 2-figure supplement 2D). Congruent with our hypothesis, NFATc2 inhibition suppressed anchorage independent growth (figure 2-figure supplement 2E), while ectopically expressed NFATc2 facilitated colony formation (figure 2-figure supplement 2F). *In vivo* subcutaneous xenograft models showed NFATc2 knockdown significantly retarded tumor sizes and growth rates for HCC827 and PDCL#24, respectively (Fig. 2H, figure 2-figure supplement 3A), while NFATc2 overexpression in A549 led to the reciprocal effects (Fig. 2I). To evaluate TIC frequencies *in vivo,* limiting dilution assays were performed by subcutaneous transplantation of decreasing numbers of tumor cells in nude mice. NFATc2 knockdown led to reduction in tumor incidence with significantly reduced TIC frequency of HCC827 xenografts (Fig. 2J, figure 2-figure supplement 3B). Reciprocal effects were observed with NFATc2-overexpression, where the TIC frequency of A549 xenografts was significantly increased (Fig. 2K, figure 2-figure supplement 3C). Together, the data supported NFATc2 mediates *in vitro* and *in vivo* TIC properties.

### NFATc2 promoted cancer resistance to cytotoxic and targeted therapy

The effect of NFATc2 on treatment response was first investigated using cisplatin chemotherapy on PDCL#24, a *KRAS V12D* mutant lung AD cell line raised from a local male chronic smoker. Upon NFATc2 inhibition, the drug response was sensitized with significantly reduced IC_50_ compared to control cells (Fig. 3A). Similarly for HCC827, NFATc2 knockout significantly reduced cisplatin IC_50_ (Figure 3-figure supplement 1A). In contrast, overexpressing NFATc2 in A549 increased cisplatin resistance with significantly elevated IC_50_ (Fig. 3B). *In vivo*, mice bearing PDCL#24 xenografts treated with cisplatin alone showed 1.68 fold tumor shrinkage compared with vehicle control. With additional stable NFATc2 knockdown, tumor shrinkage was significantly accentuated to 7 and 8.6 folds (p<0.01), respectively (Fig. 3C), while tumor growth rate was also retarded (Figure 3-figure supplement 1B). Besides, NFATc2 knockdown significantly increased paclitaxel sensitivity of HCC827 cells (Figure 3-figure supplement 2A). Furthermore, chronic exposure to increasing doses of cisplatin was used to induce resistant A549 cancer cells (A549CR). This led to NFATc2 upregulation with elevated IC_50_ compared to parental A549 cells (Fig. 3D-E). With NFATc2 suppression, A549CR was re-sensitized with return of IC_50_ to around the pre-induction level (p<0.01) (Fig. 3E). In line with this observation, induction of H1299 for paclitaxel resistance also caused NFATc2 upregulation (Figure 3-figure supplement 2B).

**Figure 3.**
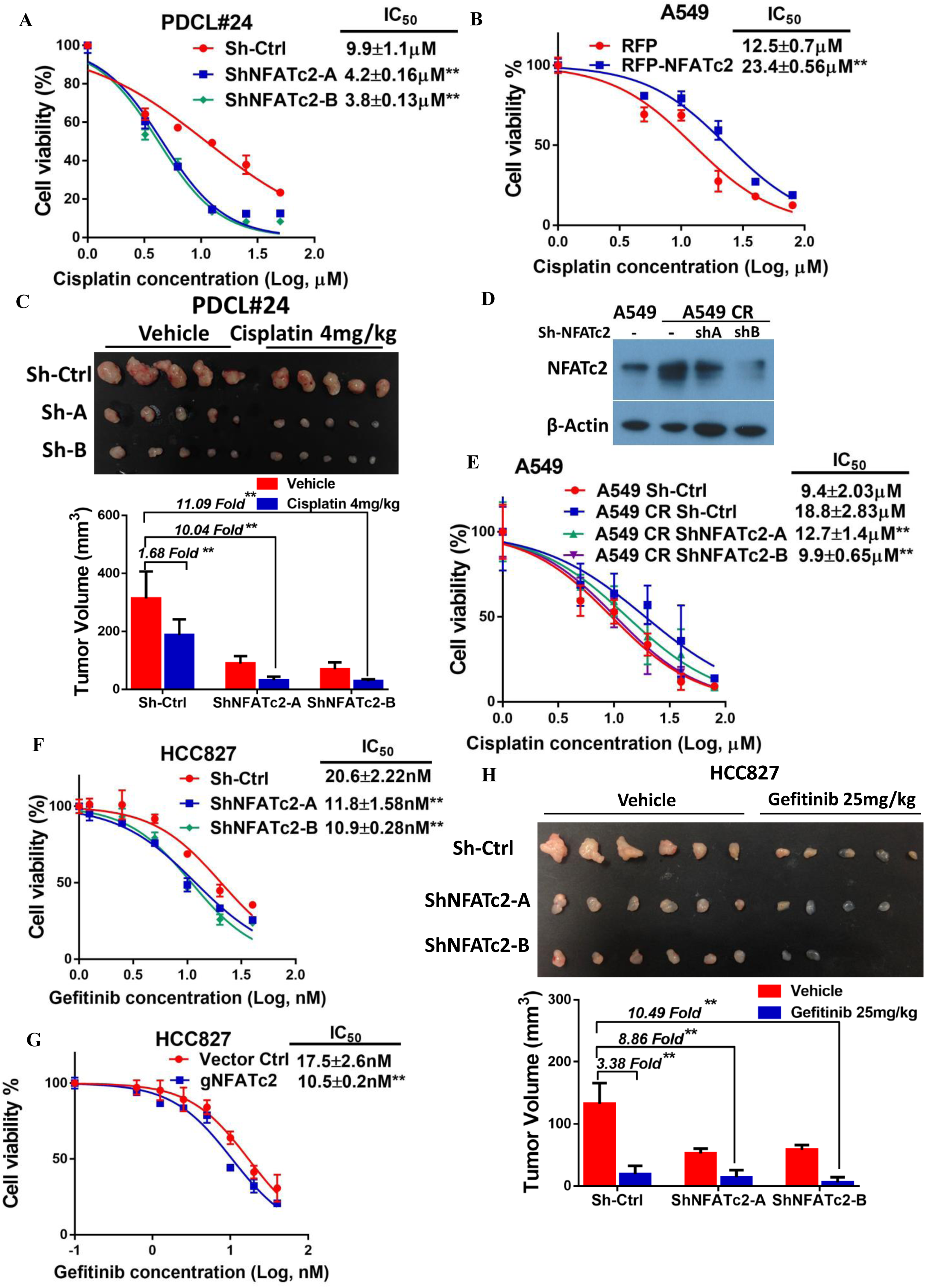
NFATc2 promoted cancer resistance to cytotoxic and targeted therapy. **(A-B)** Cisplatin response analyzed by MTT assay in PDCL#24 cells with NFATc2 knockdown (A) and A549 cells with NFATc2 overexpression (B). *p<0.05, **p<0.01 versus control by t-test. Error bar indicates the mean ± SD for at least three independent replicates. **(C)** *In vivo* cisplatin response analysis of PDCL#24 xenografts. Nude mice bearing subcutaneous xenografts were randomly separated into two groups and treated with intraperitoneal injections of cisplatin (4mg/kg every three days) or saline control, respectively. Xenografts were photographed and histograms of tumor volumes were compared to vector and no-treatment controls. **p<0.0001 versus control vehicle by two-way ANOVA. Error bar indicates the mean ± SD of tumor volumes of five mice for Sh-Ctrl vehicle and Sh-NFATc2-A vehicle groups (1 mouse from each group failed to develop tumor) and six mice for other groups. (**D-E)** Expression of NFATc2 analyzed by Western blot (D), and cisplatin sensitivity analyzed by MTT assay (E) in A549 and A549 CR cells with or without NFATc2 knockdown. **(F-G)** MTT assay for HCC827 cells with NFATc2 knockdown (F), or knockout (G), treated with gefitinib for 72 hrs. E-G: **p<0.01 versus control by t-test. Error bar indicates the mean ± SD for at least three independent replicates. **(H)** Effects of NFATc2 stable knockdown on *in vivo* response of HC827 xenografts to gefitinib. Nude mice bearing subcutaneous xenografts were randomly separated into two groups and treated with gefitinib (25mg/kg/day by oral gavage) or 1% Tween 80 as control. Xenografts were harvested. **p<0.0001 versus control vehicle by two-way ANOVA. Error bar indicates the mean ± SD of tumor volumes of six mice. The following figure supplements and source data are available for figure 3: Figure 3-figure supplement 1: NFATc2 promoted cancer resistance to cisplatin treatment. Figure 3-figure supplement 2: NFATc2 promoted cancer resistance to paclitaxel treatment. Figure 3-figure supplement 3: NFATc2 promoted cancer resistance to gefitinib treatment Figure 3-source data 1: Statistical analyses for figure 3C and H.

HCC827 is a lung AD cell line known to harbor an activating *EGFR* exon19 deletion which sensitizes it to tyrosine kinase inhibitor (TKI) therapy. To investigate whether NFATc2 contributes to resistance to targeted therapy, NFATc2 was stably inhibited by shRNA knockdown or CRISPR knockout, which led to significantly reduced IC_50_ for gefitinib (Fig. 3F, G). *In vivo,* while either gefitinib treatment or NFATc2 knockdown alone was associated with smaller HCC827 xenografts and slower growth rates compared to their respective controls, the combined treatment accentuated the level of xenograft shrinkage (Fig. 3H and figure 3-figure supplement 3A). Further, HCC827 induced for gefitinib resistance (HCC827GR) showed upregulated NFATc2 and increased NFAT promoter activities compared to parental cells (Figure 3-figure supplement 3B and C). With NFAT inhibition by the calcineurin/NFAT inhibitor cyclosporin A (CSA), drug sensitization with significantly reduced IC_50_ for gefitinib was effected (Figure 3-figure supplement 3D).

Integrating the *in vitro* and *in vivo* data of various combinations of multiple cell lines and cancer drug treatments, the enhancing effect of NFATc2 on drug resistance to cytotoxic and targeted therapy was demonstrated.

### NFATc2 upregulated SOX2 expression through its 3’ enhancer

To understand the molecular mechanism through which NFATc2 mediates cancer cell stemness and drug resistance, we hypothesize NFATc2 might be linked to the pluripotency machinery through its regulatory action. Indeed, analysis of 4 lung AD cell lines showed transcripts of the major stemness factors *SOX2*, *OCT4* and *NANOG* were significantly elevated in tumorspheres compared to monolayers (Figure 4-figure supplement 1). Genetic inhibition of NFATc2 in HCC827 (Fig 4A) and PDCL#24 (Figure 4-figure supplement 2A) led to consistent *SOX2* repression with the highest magnitude of change compared to the other 2 factors (p<0.01), while all 3 were significantly upregulated on NFATc2 ectopic expression (Fig. 4B, Figure 4-figure supplement 2B). Corresponding changes were shown at the protein level (Fig. 3C, Figure 4-figure supplement 2C). Together, the data suggested SOX2 is a major stemness target of NFATc2.

**Figure 4.**
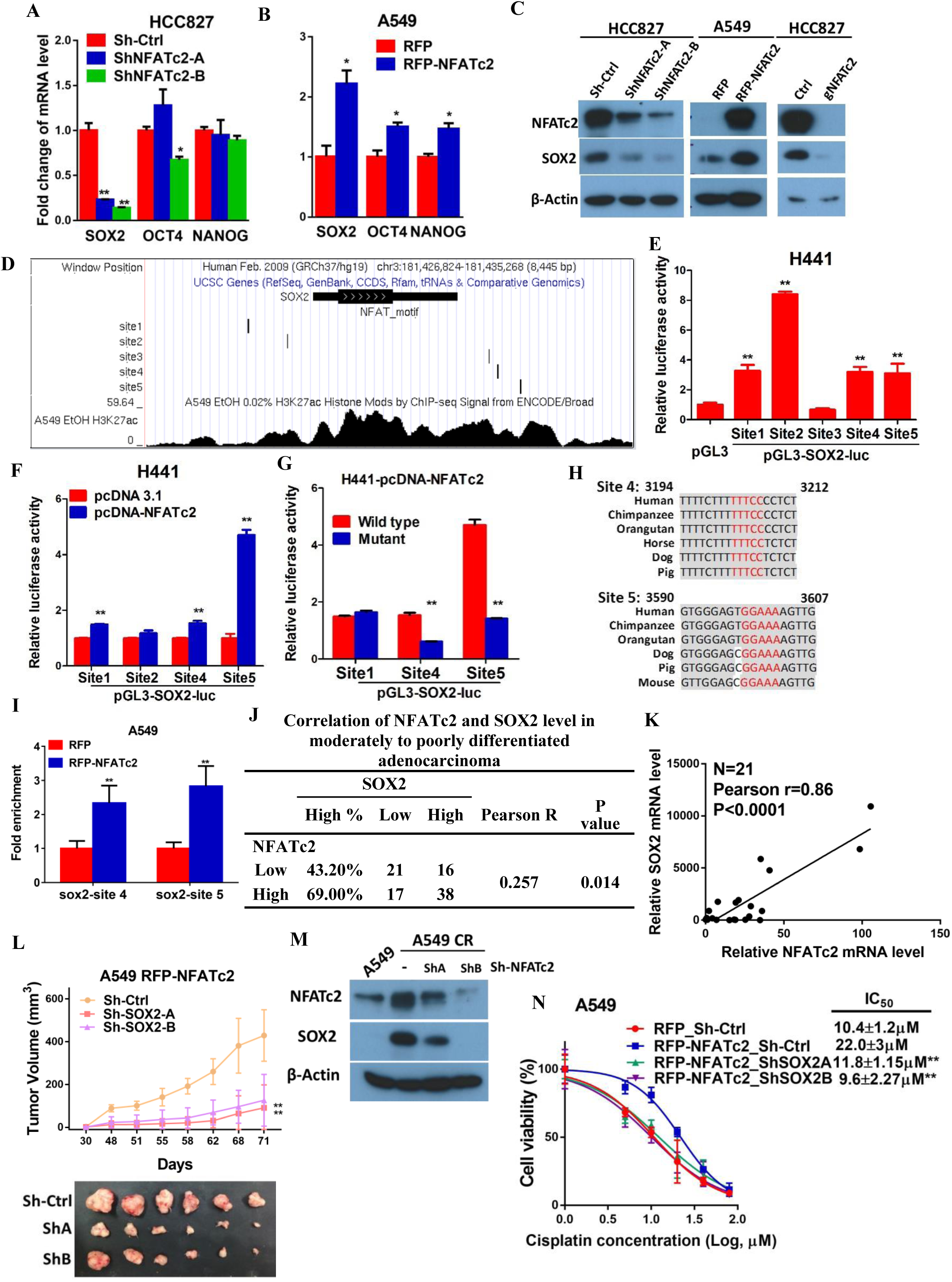
NFATc2 regulated tumor function through trans-activating SOX2 expression. **(A-B)** Pluripotency genes expressions analyzed by qPCR indicated cells. (**C)** Effects of stable NFATc2 knock-down, overexpression, or knockout on SOX2 expression in lung cancer cells by Western blot analysis. **(D)** Genome browser view of NFAT binding sites (site 1 to 5 shown as black curve) on SOX2 regulatory region 2 and 3 with H3K27Ac mark in A549 cells (shown in black). **(E)** Promoter activities of site 1-5 by dual luciferase reporter assays in H441 cells. **(F)** Effect of transient NFATc2 over-expression on transcriptional activities of indicated reporters in H441 cells. **(G)** Site-directed mutagenesis of NFAT binding sequences in indicated NFAT reporters was performed. Reporter activity of wild type and the corresponding mutant reporters were studied in H441 cells with transient NFATc2 overexpression. **(H)** Sequence alignment of site 4 and site 5 sequences in different species. Putative NFAT binding sites are highlighted in rectangle. Identical sequences were highlighted in gray. **(I)** Confirmation of NFATc2 binding to SOX2 sites by ChIP–qPCR analysis in A549 cells with stable NFATc2 overexpression. **(J)** Correlation of immunohistochemical expressions of NFATc2 and SOX2 in 92 moderately to poorly differentiated human lung adenocarcinoma by χ^2^-test. Pearson r: Pearson correlation coefficient. **(L)** *In vivo* tumorigenicity of A549 cells with NFATc2 overexpression and SOX2 knockdown was assessed by subcutaneous inoculation of 1×10^4^ cells into SCID. Xenograft formation was monitored by tumor growth curves and tumor sizes. **p<0.0001 versus control vehicle by two-way ANOVA. Error bar indicates the mean ± SD of tumor volumes of six mice. **(M)** Effect of NFATc2 knockdown on SOX2 expression in A549 CR cells analyzed by immunoblot. **(N)** MTT assay of cisplatin sensitivity for NFATc2-overexpressing A549 cells with stable knockdown of SOX2. For A, B, E-G, I, N *p<0.05, **p<0.01 versus control by t-test. Error bar indicates the mean ± S.D. for at least three independent replicates. The following figure supplements and source data are available for figure 4: Figure 4-figure supplement 1: Expression of pluripotency factors in tumorspheres. Figure 4-figure supplement 2: NFATc2 regulated SOX2 expression Figure 4-figure supplement 3: NFATc2 regulated SOX2 expression through binding to 3’ regulatory regions. Figure 4-figure supplement 4: NFATc2 regulated tumor function through SOX2. Figure 4-source data 1: Statistical analyses for figure 4K.

To further delineate the molecular mechanism of SOX2 regulation by NFATc2, we screened, *in silico,* the genomic sequences spanning 5 kb up- and downstream of the *SOX2* transcription start site (TSS), which identified 4 regions encompassing multiple conserved NFAT binding sequences (Figure 4-figure supplement 3A). Alignment with ChIP-seq data of A549 cells retrieved from public databases showed significant overlap at loci of H3K27ac occupancy with regions 2 and 3, respectively, implicating these regions might harbor sites of active transcription, which was confirmed by luciferase assay (Fig. 4D). To evaluate this possibility, SOX2 luciferase reporter assays of the putative sites were tested which revealed significant activities mediated by sites 1, 2, 4, and 5 (Figure 4-figure supplement 3B, Fig. 4E). Using H441 lung cancer cell line with transient NFATc2 overexpression, significant enhancement was observed for sites 1, 4 and 5 only (Fig. 4F). Further delineation by site directed mutagenesis of NFAT motifs (GGAAA to GACTA) prevented reporter activities of sites 4 and 5 only (Fig. 4G), and the findings were supported by data from A549 and H1299 cells bearing stable NFATc2 overexpression (Figure 4-figure supplement 3C, D). Notably, sequence homology analysis showed *SOX2* sites 4 and 5 are highly conserved across different mammalian species (Fig. 4H). Next, using NFATc2-overexpressing cell lines, NFATc2 ChIP-qPCR assays yielded significant enrichment of sites 4 and 5 sequences compared to vector (Fig. 4I), or IgG control (Figure 4-figure supplement 3E), respectively, demonstrating physical binding of NFATc2 to *SOX2.* Together, the data showed NFATc2 upregulates SOX2 by binding to its 3’ enhancer region at around 3.2kb (site 4) and 3.6kb (site 5) from the TSS, respectively.

### NFATc2 and SOX2 expressions were significantly correlated in human lung AD and NFATc2/SOX2 coupling augmented tumor functions

The clinical significance of NFATc2/SOX2 coupling was first assessed in human lung cancers. To avoid the confounding effect of *SOX2* gene amplification in squamous cell carcinoma (SCC) and focusing on tumors in which we demonstrated the role of NFATc2, IHC was performed on 92 moderately to poorly differentiated AD which showed significant correlation between SOX2 and NFATc2 expressions (Fig. 4J). In lung AD cell lines, *NFATc2* and *SOX2* transcripts expression were also positively correlated (Fig. 4K). Together, the data supported NFATc2 upregulates SOX2 in clinical and cultured lung AD.

Next, we evaluated whether NFATc2-induced SOX2 upregulation was functionally relevant for its role in sustaining TIC. Using A549 transduced for NFATc2 overexpression, SOX2 suppression by 2 shSOX2 sequences led to significantly reduced tumorspheres formation (Figure 4-figure supplement 4A, B), and cell motility (Figure 4-figure supplement 4C). *In vivo,* NFATc2-mediated enhancement of xenograft size and tumor growth rate were also abrogated by SOX2 knockdown (Fig. 4L).

Similar to NFATc2, SOX2 was upregulated in A549 induced for cisplatin resistance (A549CR) but on NFATc2 knockdown, SOX2 levels were repressed (Fig. 4M), suggesting NFATc2/SOX2 coupling was functionally active in resistant cancer cells. In MTT assays of A549 cells, while NFATc2 overexpression induced cisplatin resistance, SOX2 silencing restored sensitivity to a level comparable to the control cells (Fig. 4N). Overall, the data indicated NFATc2 induces TIC, cancer initiating phenotypes and drug resistance through upregulating SOX2 expression.

### ALDH1A1 was a target of NFATc2/SOX2 regulation

ALDH^+^/CD44^+^-TIC was investigated as a candidate cellular target of NFATc2. The ALDH^+^/CD44^+^ subset isolated by flow cytometry from HCC827 and H1650 cell lines showed significantly higher levels of *NFATc2* and *FASL* transcripts compared to ALDH^-^/CD44^-^ (Fig. 5A), suggesting NFATc2 played a role in regulating this cell population. When NFATc2 was knocked down or knocked out in HCC827, ALDH^+^/CD44^+^-TIC was significantly reduced (Fig. 5B, C). Consistent changes were observed for PDCL#24 with NFATc2 knockdown (Figure 5-figure supplement 1A). Conversely, in both A549 with NFATc2 overexpression and in A549CR cells, ALDH^+^/CD44^+^-TIC proportions were increased (Figure 5-figure supplement 1B-C). Breakdown analysis showed the trend of changes were consistent with those of the ALDH^+^ but not CD44^+^ population, suggesting ALDH might be the main target of NFATc2.

**Figure 5.**
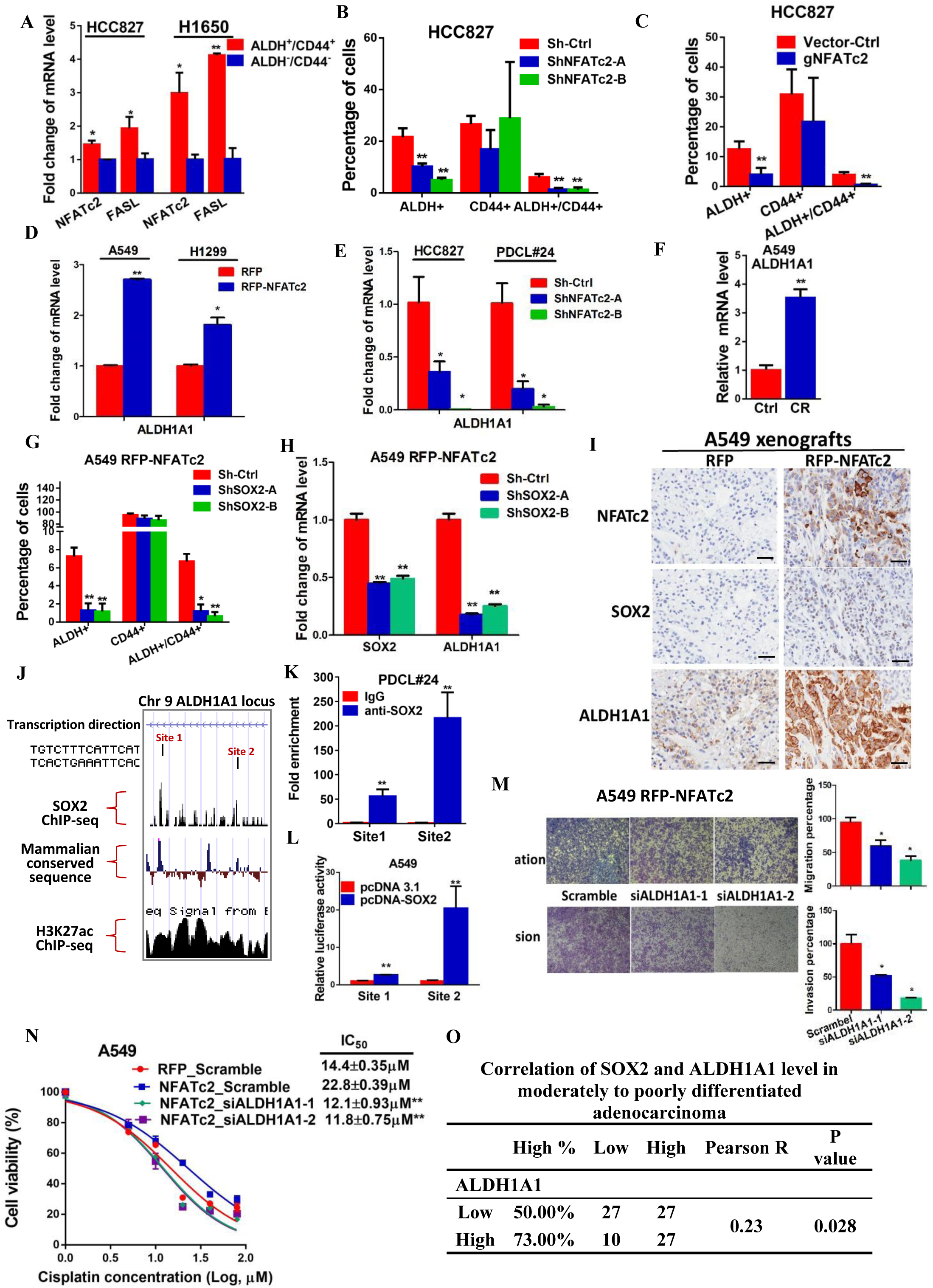
ALDH1A1 was a target of NFATc2/SOX2. **(A)** Expression of *NFATc2* and its target *FASL* analyzed by qPCR in ALDH^+^/CD44^+^ TIC population compared to ALDH^-^/CD44^-^ population. (**B-C)** Flow cytometry analysis of TIC proportions by ALDH^+^/CD44^+^ markers, in HCC827 cells with NFATc2 knockdown (B), or knockout (C). **(D-F)** mRNA levels of *ALDH1A1* analyzed by qPCR in cells with NFATc2 up-(D) or down-regulation (E), or A549 CR cells (F). **(G)** Effects of SOX2 knockdown in NFATc2-overexpressing cells on ALDH/CD44 distribution by flow cytometry. **(H)** Expression of *SOX2* and *ALDH1A1* in A549 cells with NFATc2 overexpression and SOX2 knockdown. **(I)** Representative image of immunohistochemical expression of NFATc2, SOX2 and ALDH1A1 in xenografts derived from indicated A549 cells. Scale bars, 50 µM. **(J)** ChIP-seq genome browser view of SOX2 peak with SOX2 binding motif (curved arrow) on ALDH1A1 enhancer. The SOX2 peaks were co-localized with mammalian conservation peaks (blue) and H3K27Ac peak in A549 cells (black). (**K)** Confirmation of SOX2 binding to ALDH1A1 sites by ChIP–qPCR analysis in PDCL#24 cells. **(L)** Site 1 and 2 reporters were co-transfected with SOX2 overexpressing vectors and reporter activity was analyzed by dual luciferase reporter assay in A549 cells. **(M-N)** Effects of transient ALDH1A1 suppression on invasion and migration abilities (M), and cisplatin sensitivity (N) of indicated cells. **(O)** Correlation between ALDH1A1 and SOX2 expressions by IHC in human lung adenocarcinomas by χ^2^-test. *P<0.05, **p<0.01 versus control by t-test. Error bar indicates the mean ± S.D. for at least three independent replicates. The following figure supplements and source data are available for figure 5: Figure 5-figure supplement 1: NFATc2 regulated the ALDH+ population. Figure 5-figure supplement 2: NFATc2 regulated ALDH1A1 expression. Figure 5-figure supplement 3: Effect of siALDH1A1 on ALDH1A1 expression. Figure 5-figure supplement 4: Effect of NFATc2/SOX2 on β-catenin activity. Figure 5-source data 1: Statistical analyses for figure 5B, C and G.

ALDH1A1 is the most frequent and important ALDH isozyme reported in lung cancer TIC (Ucar, Cogle et al. 2009, Tomita, Tanaka et al. 2016). Although ALDH1 is marketed as the major subtype contributing to ALDH activities detected by the ALDEFLUOR^TM^ assay, cross-reactivity with other isoforms cannot be excluded. Hence, to explore the part contributed by ALDH1A1, we abrogated ALDH1A1 in A549 engineered to overexpress NFATc2, which led to significantly suppressed aldefluor activities (Figure 5-figure supplement 2A). Furthermore, in cancer cells with NFATc2 up-or down-regulation (Fig. 5D, E), in A549CR (Fig. 5F), or upon NFATc2 knockout (Figure 5-figure supplement 2B), ALDH1A1 mRNA were correspondingly altered, indicating NFATc2 regulates ALDH1A1 expression which contributes to the majority of ALDH positivity in ALDH^+^/CD44^+^-TIC.

Further analyze of the role of NFATc2/SOX2 coupling in ALDH1A1 regulation showed silencing SOX2 in NFATc2-overexpressing A549 cells consistently prevented the increase of ALDH^+^ and ALDH^+^/CD44^+^ subpopulations only (Fig. 5G). Specifically, the expected upregulation of *ALDH1A1* was also abolished (Fig. 5H). In addition, as analyzed by IHC, xenografts derived from NFATc2-overexpressing A549 cells showed corresponding upregulation of SOX2 and ALDH1A1 (Fig. 5I), while NFATc2 suppression in PDCL#24 led to congruent changes with marked SOX2 and ALDH1A1 downregulation (Figure 5-figure supplement 2C), respectively. Together, the data strongly supported NFATc2/SOX2/ALDH1A1 form a regulatory axis in lung cancer.

To examine the mechanism of SOX2 in ALDH1A1 regulation, ChIP-seq analysis was performed to identify SOX2 binding sequences in PDCL#24 cell line. Two loci of peak signals encompassing the SOX2 consensus binding motif (ATTCA) were identified in the 5’ region of the *ALDH1A1* gene at around 27 kb from the TSS (site 1 and 2, Fig. 5J). Bioinformatics analysis showed these loci were located at chr9:75,595,820-75,595,832 and chr9:75,602,860-75,602,872 which overlap with H3K27ac occupancy deposited in public ChIP-seq database of A549 cells as well as mammalian conserved sequences (Fig 5J). On the other hand, no SOX2 binding motif was identified in regions flanking other *ALDH* isoform genes. Together, the findings suggested the presence of an accessible and highly conserved chromatin region encompassing putative SOX2 binding motifs 5’ to the *ALDH1A1* gene. To confirm, ChIP-q-PCR assay using PDCL#24 cells was performed which showed anti-SOX2 antibodies significantly enriched both sequences 1 and 2 (Fig. 5K). To study their transcriptional regulatory role, luciferase reporter constructs for sites 1 and 2 were co-transfected with SOX2 expression plasmids into A549 cells which yielded significantly enhanced reporter activities compared to control cells (Fig. 5L). Compatible results were obtained for A549 with NFATc2 overexpression (Figure 5-figure supplement 2D).

To evaluate whether ALDH1A1 is a functionally relevant target, ALDH1A1 was knocked-down by siRNA in NFATc2-overexpressing A549 cells (Figure 5-figure supplement 3). This led to significant suppression of cell motility (Fig. 5M) and cisplatin sensitization (Fig. 5N). Furthermore, moderately to poorly differentiated human lung AD showed statistically significant positive correlation between SOX2 and ALDH1A1 expressions by IHC staining (Fig. 5O). Collectively, the data supported ALDH1A1 is a functional target of regulation through NFATc2/SOX2 coupling.

### NFATc2/SOX2/ALDH1A1 coupling enhanced drug resistance and tumor properties through ROS attenuation

Alleviation of oxidative stress induced by chemotoxicity promotes cancer cell survival and mediates drug tolerance. Thus, in A549CR cells, intracellular ROS levels were significantly lower compared to parental A549 cells (Fig. 6A). To investigate for possible relation between NFATc2 and ROS modulation, NFATc2 was silenced by knockdown or knockout which led to increased ROS levels (Fig. 6B-D). As shown in Fig. 6E, NFATc2 depletion sensitized PDCL#24 cells to cisplatin treatment, but the addition of the reducing agent NAC reversed cisplatin IC_50_ to above the control level dose-dependently. Reciprocally, the enhanced resistance of A549 by NFATc2 overexpression was reversed by oxidative stress induced by the glutathione inhibitor BSO (Fig. 6F), consistent with the suggestion that drug resistance by NFATc2 is effected through ROS attenuation. Similarly, ROS regulation also supported other tumor phenotypes mediated by NFATc2. For example, tumorspheres suppression by NFATc2 knockdown was restored by NAC dose-dependently but in control cells, no significant changes were induced even in the presence of additional NAC (Fig. 6G-H). Likewise, cell migration and invasion efficiencies inhibited by NFATc2 depletion were reversed by NAC (Fig. 6I). To further address the involvement of SOX2 coupling and ALDH1A1, we showed suppression of ROS by NFATc2-overexpression in A549 cells were reversed by silencing SOX2 or ALDH1A1, respectively (Fig. 6J-K). Together, the data suggested NFATc2/SOX2/ALDH1A1 form a functional axis in the homeostatic regulation of an optimal level of ROS for *in vitro* tumorigenicity, cell motility, and mediation of drug resistance.

**Figure 6.**
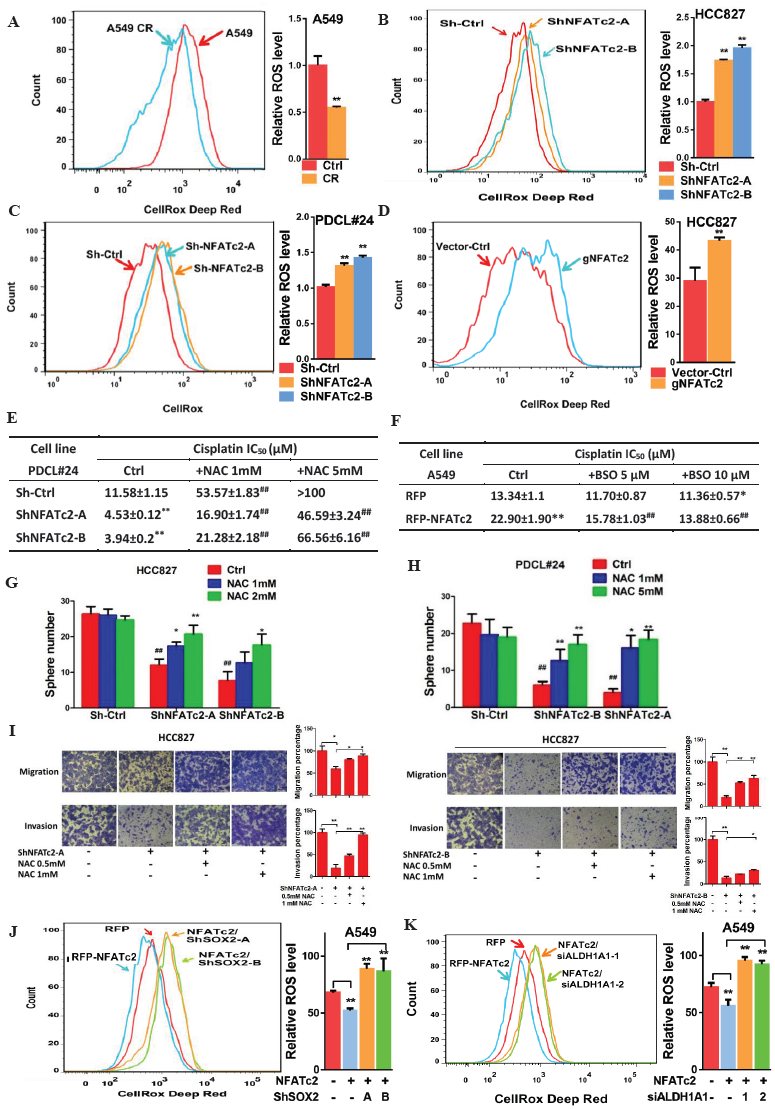
NFATc2 regulated TIC properties through ROS suppression. **(A)** ROS levels detected by flow cytometry in A549 and A549 CR cells. **(B-C)** ROS levels in HCC827 cells (B) and PDCL#24 cells (C) with or without NFATc2 stable knockdown. **(D)** ROS levels in HCC827 cells with or without NFATc2 knockout. **(E-F)** Cisplatin sensitivity expressed as IC_50_ by MTT assays of NFATc2-silenced PDCL#24 cells treated with increasing doses of NAC (E), or NFATc2-overexpressing A549 cells treated with the oxidizing agent (BSO) (F), respectively. * p<0.05, ** p<0.01 versus vector control without REDOX reagents; ^##^ p<0.01 versus the corresponding treatment control; t-test. Error bar indicates the mean ± S.D. for three independent replicates. **(G-H)** Effects of increasing doses of NAC on tumorsphere formation ability of HCC827 (G) cells and PDCL#24 cells (H). * p<0.05, ** p<0.01 versus corresponding treatment controls, ^##^ p<0.01 versus vector control, by t-test. Error bar indicates the mean ± S.D. for three independent replicates. **(I)** Effects of increasing doses of NAC on cell migration and invasion ability of HCC827 cells with NFATc2 knockdown. **(J-K)** ROS levels in NFATc2 overexpressing A549 cells with stable SOX2 (J) or transient ALDH1A1 (K) knockdown. For A-D, I-K *p<0.05, **p<0.01 versus respective control by t-test. Error bar indicates the mean ± S.D. for at least three independent replicates. Figure 6-source data 1: Statistical analyses for figure 5A-D, J and K.

## Discussion

The elucidation and disruption of TIC maintenance pathways offer the opportunity to eliminate the most resilient cancer cells and improve treatment outcome. Many studies have demonstrated cell populations expressing high levels of specific markers such as ALDH, CD44, CD166, CD133, etc. are enhanced for a multitude of tumor phenotypes, with tumorigenicity and drug resistance being clinically the most important. Constitutive stem cell programs or stress-induced pathways are the main TIC sustaining mechanisms but details of their regulation are still elusive. NFAT is a family of transcription factors with the calcium-responsive isoforms NFATc1, -c2, -c3 and -c4 being expressed in a tissue-dependent manner. In both clinical NSCLC and lung AD cell lines, we observed significantly upregulated transcripts of NFATc2, and -c4 compared to paired normal lung and immortalized bronchial epithelial cells BEAS-2B, respectively, while NFATc2 was the most frequently involved entity with the highest magnitude of change (data not shown). In this study, using NFATc2 depletion and overexpression models of multiple lung cancer cell lines in TIC-defining functional assays (Pattabiraman and Weinberg 2014), as well as clinical evidence from excised human lung cancers, we showed NFATc2 mediates TIC phenotypes. *In vitro* cell renewability was demonstrated by tumorspheres passaged for consecutive generations and augmentation of tumorigenicity was illustrated by the limiting dilution assay. In clinical tumors, high level NFATc2 segregated with impaired tumor differentiation, advanced pathological stage, shorter recurrence-free and overall survivals in NFATc2-positive NSCLC, suggesting NFATc2 mediates the more primitive and aggressive tumor phenotypes. In the literature, only one research group has reported NFATc2 expression in 52% of 159 lung cancers and similar to our findings, high expression was associated with late tumor stage and poor survival. Supportive evidences for cell proliferation, invasion and migration were demonstrated by cell models but the *in vivo* role of NFATc2 and, in particular, its effects on TIC, drug response or mechanisms of action were not addressed (Chen, Zhao et al. 2011, Liu, Zhao et al. 2013).

To identify the TIC sustaining mechanism of NFATc2, we hypothesized NFATc2 might be coupled to the core pluripotency factors SOX2, NANOG and/or OCT4, aberrant activities of which would be most suitable to orchestrate multifaceted cancer propensities through extensive transcriptional and epigenetic reprograming. Using multiple analyses of cancer cells and tumorspheres, we observed SOX2 was the most consistently altered factor with the highest magnitude of change when NFATc2 expression was genetically manipulated. SOX2 is an important oncogene for squamous cell carcinomas (SCC) of the lung and other organs (Lu, Futtner et al. 2010, Boumahdi, Driessens et al. 2014), and aberrant expression is mostly due to amplification of the SOX2 locus at 3q26.3 (Hussenet, Dali et al. 2010). For lung AD, although SOX2 is often expressed at high levels and predicts adverse survivals (Chou, Lee et al. 2013), distinct genetic mechanisms are not identified, indicating regulatory or signaling aberrations might be involved. To evaluate the effects of NFATc2 on SOX2 expression, we avoided the potentially confounding element of SCC and focused on moderate to poorly differentiated AD where NFATc2 is shown to play an important prognostic role. Indeed, SOX2 expression is significantly correlated with that of NFATc2 in this group of human lung cancer. Functionally, NFATc2/SOX2 coupling contributes to tumor behavior as depletion of SOX2 in cell lines with NFATc2 overexpression led to significant suppression of TIC phenotypes. Hence, clinical and experimental evidences support NFATc2 impedes tumor differentiation and negatively affects patient outcome through coupling to SOX2 in lung AD.

We observed NFATc2 binds to the 3’ enhancer region of *SOX2* at around 3.2 kb and 3.6 kb from TSS, effecting functional TIC enhancement and supporting the direct involvement of NFATc2 in stemness induction. In a recent study of a transgenic mouse model of pancreatic ductal adenocarcinoma with *KRAS*^*G12VD*^ mutation and p53 heterozygous inactivation, Singh *et al* reported an analogous mechanism involving another NFAT family protein, NFATc1, which acts as a coactivator and transcriptional regulator of SOX2 (Singh, Chen et al. 2015). They showed NFATc1 played a permissive role for tumor dedifferentiation and expression of EMT genes, while p53 disruption was essential for tumorigenesis, suggesting under the appropriate genetic context, NFATc1 activation transduces EMT through SOX2 upregulation. The authors proposed since chronic inflammation is a known etiology of pancreatic AD, NFATc1 might be a crucial factor involved in the progression of this cancer. NFAT family members are distributed in a tissue-specific manner and we have observed NFATc2, rather than NFATc1, is the main factor highly expressed in lung cancers. Our data also distinguish a different SOX2 enhancer region accessible to NFATc2 which functions not only in the presence of *KRAS* mutation (A549, PDCL#24) but also in *EGFR* mutant (HCC827) cancers. Chronic inflammation is an important etiological mechanism of lung cancer through release of ROS and free radicals from alveolar macrophages and neutrophils. While many studies modeling carcinogenetic mechanisms of tobacco toxicity or chronic obstructive pulmonary diseases have featured the NF-κB pathway as a major mediator, our findings on the effects of NFATc2 on TIC induction might add to this repertoire. In fact, NFATc2 and NF-κB share highly similar DNA binding domains but differ in the upstream activators, with NF-κB being stimulated by cytokine receptors and inflammatory molecules while NFATc2 is downstream of calcium signaling reputed as a stress response integrator, illustrating the multiple mechanisms through which inflammation-induced carcinogenesis might be initiated in the lung.

We have shown the ALDH^+^/CD44^+^ fraction of lung cancer cells, despite being the smallest subset, demonstrates the highest tumorigenic capacity compared to their counterpart cell compartments (Liu, Xiao et al. 2013). In trying to understand the role of NFATc2/SOX2 coupling in inducing this cell fraction, we observed only ALDH but not CD44 showed consistent changes upon NFATc2 suppression or overexpression, respectively, and more specifically, ALDH1A1 was identified as a major functional target. Recent reports have shown β-catenin can directly regulate ALDH1A1 (Condello, Morgan et al. 2015), and since SOX2 can upregulate β-catenin (Yang, Hui et al. 2014), this could infer SOX2 might indirectly regulate ALDH1A1 through β-catenin. We thus assessed β-catenin activity by western blot in A549 cells with NFATc2 overexpression and SOX2 knockdown, but no significant alternation of total or activated β-catenin levels was observed in these cells and the possible mechanism of indirect ALDH1A1 upregulation through β-catenin was not supported (Figure 5-figure supplement 4). On the other hand, computational screening did not detect significant and conserved SOX2 binding motifs within the proximal promoter region of ALDH1A1, but a more distant locus at 27 kb upstream from its TSS was shown to be a probable response region which was identified and confirmed by CHIP-seq and luciferase reporter assays, respectively. This illustrates a distal enhancer is involved in the trans-regulation of ALDH1A1 expression, which has not been reported before. Current views on cancer stem cells suggest TIC is unlikely to be a single population with unique identifiers; instead, cell plasticity might induce variant TIC populations through dynamic response mechanisms, enabling cancer cells to meet the requirements of a complex micro-environment. In this connection, since we have only demonstrated the NFATc2/SOX2/ALDH1A1 axis, further investigation for the mechanisms regulating CD44 in ALDH^+^/CD44^+^-TIC is needed.

Acquired drug resistance is mediated through complex genetic and molecular mechanisms (Holohan, Van Schaeybroeck et al. 2013). We have shown NFATc2 augments resistance to both cytotoxic chemotherapy and targeted therapy. Adaptive antioxidant response for alleviating oxidative stress from ROS surge during systemic therapy is one of the most important mechanisms of drug resistance (Zhang, Du et al. 2011), suggested to be accentuated in cancer stem cells (Diehn, Cho et al. 2009, Achuthan, Santhoshkumar et al. 2011, Ishimoto, Nagano et al. 2011, Chang, Chen et al. 2014). Indeed, we have observed in multiple cell lines with induced resistance to chemotherapy or targeted therapy, NFATc2 was upregulated, while ROS was maintained at a lower level in A549CR compared to parental cells. Changes in TIC phenotypes induced by NFATc2 up- or down-regulation were correspondingly restored by the redox reagents BSO or NAC, respectively, suggesting ROS scavenging is an important mechanism of drug resistance and other TIC properties mediated by NFATc2. In line with this suggestion, it has been reported in adult immortalized bronchial epithelial cells, NFAT can be upregulated in response to inflammatory and carcinogenic stimulation such as those due to benzo-(a)-pyrene and heavy metals stimulation (Huang, Li et al. 2001, Ding, Wu et al. 2007, Cai, Li et al. 2011), leading to ROS-induced COX2 pathway signaling and enhanced cell survival (Ding, Li et al. 2006). However, whether ROS is in turn suppressed by NFATc2 through a negative feedback mechanism has not been reported. Our findings supplement this information and show NFATc2 facilitates ROS scavenging, and further implicates this effect is mediated by ALDH1A1 through SOX2 coupling, which is consistent with other studies showing that ALDH1A1 is involved in mediating drug resistance through repressing ROS level (Singh, Brocker et al. 2013, Raha, Wilson et al. 2014, Mizuno, Suzuki et al. 2015).

In summary, this study demonstrates the calcium signaling molecule NFATc2 enhances functional characteristics associated with cancer stemness phenotype. Our data reveal a novel mechanism of SOX2 upregulation in lung cancers through enhancer binding by NFATc2. The NFATc2/SOX2/ALDH1A1 axis contributes to drug resistance by mediating a negative feedback mechanism for ROS scavenging and restoration of redox homeostasis. Together, the findings implicate NFATc2 is a potential therapeutic target for sequential or combination therapy of lung cancer that aims to eliminate TIC.

## Materials and Methods

### Cell lines

Established cell lines (H1993, HCC1833, H358, H1650, H2228, H1299, H1437, H1975, H23, H2122, HCC827, HCC78, A549, H441, and BEAS-2B) were obtained from ATCC. HCC366 and HCC78 were kindly provided by Dr. J. Minna (University of Texas Southwestern Medical Center. Dallas). All cell lines were cultured into individual aliquots and frozen upon receipt. Only first 20 passage of cell lines were used in experiment. Patient derived cell lines (HKULC1, HKULC2, HKULC3, HKULC4, PDCL#24, and FA31) were raised from resected lung cancers or malignant pleural effusions and only cells of the 1st to 10th passage were used for study. Cancer cells were maintained in RPMI-1640 (Invitrogen, Carlsbad, CA) with 10% FBS (Invitrogen, Carlsbad, CA). BEAS-2B were cultured in Keratinocyte-SFM (Invitrogen, Carlsbad, CA). Gefitinib, paclitaxel or cisplatin-resistant (-GR, TR or –CR, respectively) cells were generated by chronic exposure of cancer cells to stepwise increasing doses of the respective drugs. All procured cell lines used in this study were authenticated using the AmpFlSTR® Identifiler® PCR Amplification Kit for short tandem repeat profiling according to the manufacturer’s instruction (Thermo Fisher Scientific, Waltham, MA).

### SiRNA and Plasmids

Small interfering RNA (siRNA) with pre-designed sequences targeting human PPP3R1, ALDH1A1 and scramble siRNA were from Sigma-Aldrich (St Louis, MO). GFP-VIVIT (11106), pGL3-NFAT luciferase (17870), two shRNA sequences targeting SOX2, pLKO.1 Sox2 3HM a (26353) and pLKO.1 Sox2 3H b (26352), the negative control vector pLKO.1-puro (1864), the envelope vector pMD2.G (12259) and packaging vector psPAX2 (12260) were purchased from Addgene (Cambrige, MA; http://www.addgene.org). The pLKO.1-lentiviral shRNA with different inserts specifically targeting NFATc2 were purchased from Sigma-Aldrich (TRCN0000016144, TRCN0000230218). Human full length NFATc2 were amplified by PCR, and the RFP-NFTAc2 plasmids were generated by cloning the sequences into PCDH-CMV-MCS-EF1-COPRFP vector (SBI, Mountain View, CA). For luciferase reporter construction, SOX2 regulatory regions were amplified by PCR from human genomic DNA and cloned into pGL3 (Promega) to generate the SOX2-luc constructs. Primers used for genomic DNA amplification were listed in supplementary table 1. Site directed mutagenesis of the consensus NFAT binding site (GGAAA to GACTA) were performed using QuikChange (Stratagene).

### Lentiviral knockdown of NFATc2 and SOX2

Lentiviral shRNA was produced by transfecting the shRNA, envelope and packaging vectors into 293T cells using lipofectamine 2000 (Invitrogen, Carlsbad, CA). Viruses were harvested after 48 hrs of transfection followed by infection of target cells for 72 hrs. Cells stably expressing shRNA were selected using puromycin (Sigma-Aldrich) for 14 days after 72 hrs of viral infection.

### Lentiviral over-expression of NFATc2

RFP-NFATc2 lentiviral particles were produced and transduced into target cells using Lenti Starter kit (SBI, Mountain View, CA) according to manufacturer’s instructions. RFP-positive cells stably over-expressing NFATc2 and SOX2 were selected by FACS using BD Aria (BD Biosciences).

### Lentiviral knock out of NFATc2 by CRISPR/Cas9

LentiCas9-Blast and lentiGuide-Puro were purchased from Addgene (Cambrige, MA; http://www.addgene.org). The gRNA targeting NFATc2 was designed using Zifit (http://zifit.partners.org/ZiFiT/) and listed in supplementary table 1. The annealed gNFATc2 oligonucleotides were cloned into lentiGuide-Puro. Lenti-viral cas9 and lenti-viral gNFATc2 were generated by transfecting lentiCas9-Blast or lenti-viral gNFATc2 together with pMD2.G and psPAX2, respectively, into 293FT cells by lipofectamine 2000 according to protocols as described (Sanjana, Shalem et al. 2014). After infection of lenti-viral cas9, cells stably expressing Cas9 were selected using Blasticidin (Sigma-Aldrich) for 10 days. HCC827-Cas9 cells were further infected with lenti-gNFATc2 virus for 72 hrs, and cells stably expressing gNFATc2 were selected using puromycin (Sigma-Aldrich) for 14 days.

### Flow cytometry and fluorescence activated cell sorting (FACS)

ALDH activity was analyzed by the Aldefluor kit (Stem Cell Technologies) according to manufacturer’s instructions. CD44 expression was stained by anti-CD44-APC (BD Pharmingen) as previously described (Liu, Xiao et al. 2013). Flow cytometry was performed using FACS Canto II (BD Biosciences) and data were analyzed using FlowJo (Tree star). RFP positive cells with NFATc2 over-expression were isolated by FACS using BD Aria (BD Biosciences). Sorted cells were re-analyzed after collection to ensure a purity of > 95%. Non-viable cells were identified by propidium iodide inclusion.

### Sphere formation and serial passage

Five hundred cells were seeded in an ultra-low plate (Costar) and cultured in cancer stem cell medium (RPMI-1640 medium supplemented with 20 ng/mL FGF, 20 ng/mL EGF, 40 ng/mL IGF and 1X B27 (Invitrogen, Carlsbad, CA) for 14 days. Tumorspheres were harvested, dissociated with trypsin, re-suspended in RPMI-1640, and 500 cells were seeded again under previously described stem cell culture conditions for generation of 2^nd^ passage.

### Colony formation assay

Cells were seeded into 6-well plates at a density of 500 cells per well. After culturing for 10 to 14 days, cells were fixed with methanol and stained with crystal violet. Colonies comprising >50 cells were counted.

### Anchorage - independent growth assay

Six-well plates coated with a layer of 0.5% agar dissolved in RMPI-1640 medium with 10% FBS were used for plating of 2,500 cells suspended in RMPI-1640 with 0.35% agarose. After 4 weeks, the soft agar was stained with crystal violet, and colony numbers were determined under light microscopy.

### Cell motility assessment by migration and invasion assay

The migration and invasion assays were performed using Corning® Transwell. Both chambers were filled with RMPI-1640 medium, and the lower chamber was supplemented with 10% FBS. For the migration assay, 5×10^4^ cells were seeded into the upper chamber and allowed to migrate for 24 hrs. For the invasion assay, the upper chamber was first coated with Matrigel (BD Pharmingen); 1×10^5^ cells were seeded and allowed to invade for 24 to 36 hrs. Cells that migrated or invaded to the lower surface of the transwells were fixed with methanol and stained with crystal violet. Cell densities were photographically captured in three random fields. The dye on the transwell membrane was dissolved by 10% acetic acid, transferred to a 96 well plate, and the dye intensity was measured by a plate spectrophotometer at 570 nm.

### Drug sensitivity assays

Drug sensitivity was tested by MTT assays. 6000 cells per well were seeded into 96-well plates and incubated for 24 hrs at 37°C, followed by exposure to gefitinib (Selleckchem Houston, TX), paclitaxel (Sigma-Aldrich, St Louis, MO), or cisplatin (Sigma-Aldrich, St Louis, MO) at various concentrations for 72 hrs with or without CSA (Selleckchem, Houston, TX), NAC or BSO (Sigma-Aldrich, St Louis, MO). Subsequently, Thiazolyl Blue Tetrazolium Bromide (MTT) (Sigma-Aldrich, St Louis, MO) was added and the mixture was incubated at 37°C for 4 hrs. The absorbance was read at 570 nm using a plate spectrophotometer. The drug response curve was plotted and IC_50_ was calculated using nonlinear regression model by GraphPad Prism 7.0.

### Quantitative PCR (qPCR) analysis

Total RNA was isolated using RNAiso Plus reagent (Takara, Mountain View, CA) and complementary DNA (cDNA) was generated using PrimeScript RT Reagent Kit (Takara, Mountain View, CA) according to the manufacturer’s instructions. Gene mRNA levels were analyzed by quantitative RT-PCR (qPCR) (7900HT, Applied Biosystems, Carlsbad, CA) and SYBR green (Qiagen, Hilden, Germany) detection. Average expression levels of *RPL13A* and *beta-2-microglobulin* (*B2M*) were used as internal controls. Primers were listed in supplementary table 2.

### Western blot analysis

Cells were harvested and lysed on ice by lysis buffer [50mM Tris HCl pH 7.4, 1% Triton X-100, 1mM EDTA, 150 mM NaCl, 0.1% SDS, with freshly added 1:50 Phosphatase Inhibitor Cocktail 2 (Sigma), 1:50 Protease Inhibitor Cocktail (Sigma)] for 30 min. The cell lysate was then centrifuged at 13k rpm for 20 min at 4°C to remove cell debris. The protein amount was quantified by the Dc Protein Assay (Bio-Rad). Cell lysates were resolved by 6-10% SDS-PAGE and then transferred onto PVDF membranes (Millipore). Primary antibodies including SOX2 (1:1000), NFATc2 (1:1000), β-catenin (1:1000), non p-β-catenin (1:1000), p-βcatenin (1:1000) or ACTIN (1:1000) (Cell Signaling, Beverly, MA), respectively, where appropriate, were added. After overnight incubation, the membrane was washed with PBS and then incubated with the anti-rabbit secondary antibody. Target proteins on the membrane were visualized on X-ray films using ECL Plus Western Blotting Detection Reagents (Amersham, Buckinghamshire, UK).

### Chromatin immunoprecipitation (ChIP)-qPCR assay

ChIP assay was performed using the Magna ChIP^TM^ A kit (Millipore, Billerica, MA) according to manufacturer’s instructions. Briefly, cells were sonicated and lysed after protein/DNA cross-linking by 1% formaldehyde for 10 min. The crosslinked complex was immuno-precipitated by anti-NFATc2 antibody or normal rabbit IgG (Cell Signaling, Beverly, MA) bound to protein A magnetic beads. After overnight incubation at 4°C, the complex was eluted and DNA was purified. The immune-precipitated DNA was quantified by qPCR using primer sequences designed to detect specific regulatory regions listed in Supplementary Table S2.

### ChIP-seq assay

ChIP assay was performed using the EZ-Magna ChIP™ A/G Chromatin Immunoprecipitation Kit (Millipore, 17-10086) according to manufacturer’s instructions. Cells were cultivated and treated with 1% formaldehyde to crosslink the protein and DNA. cell lysate was sonicated to reduce the DNA length from 100 to 500 bp. The DNA-protein fragments were then incubated with 10ug SOX2 antibodies (Abcam) and magnetic beads coated with protein A/G to form DNA-protein-antibody complex. The DNA was isolated and purified by Spin column and sent to the company(BGI) for library construction and sequencing using Illumina Hi-Seq platforms. Sequence reads were aligned to Human Reference Genome (hg19) using Bowtie (Langmead, Trapnell et al. 2009). Model based analysis of ChIP-Seq(MACS) was used for peaks identification by comparing ChIP sample over input sample with default parameters (Zhang, Liu et al. 2008).

### NFATc2 binding sites predication

The 5’- and 3’-flanking regions (-5000 to +5000 bp) of SOX2 were scanned for NFAT binding sequences using PWMSCAN (Levy and Hannenhalli 2002). The significance of the predicted sites was evaluated statistically using a permutation-based method and comparison with occurrence of the motif in background genomic sequences of intergenic regions. Phylogenetically non-conserved binding sites were filtered (Li, Sham et al. 2010).

### Luciferase reporter assay

Cells were transfected with luciferase reporters, expression plasmids and pRL-TK vector using lipofectamine 2000 (Invitrogen, Carlsbad, CA). Luciferase activity were measured by using the Dual-Luciferase Reporter Assay System (Promega).

### *In vivo* tumorigenicity

All animal experiments were performed after approval by the Animal Ethics Committee, the University of Hong Kong according to issued guidelines. Briefly, different numbers of cells mixed with an equal volume of matrigel (BD Pharmingen) were injected subcutaneously at the back of 6-week old severe combined immunodeficiency (SCID) mice or Ncr-nu/nu-nude mice. Tumors sizes were monitored every 3 days using digital vernier calipers, and tumor volumes were calculated using the formula [sagittal dimension (mm) × cross dimension (mm)^2^] / 2 and expressed in mm^3^.

### Reactive Oxygen species (ROS) measurement

Cells with or without respective treatments were washed with PBS and stained with 1 µM the ROS probe CellROX^TM^ Deep Red (Lift Technologies) for 30 mins according to manufacturer’s instructions. Fluorescence was measured by flow cytometry (FACSCanto II Analyzer, BD Biosciences) and data were analyzed using FlowJo (Tree star).

### Human lung cancers

Surgically resected primary human NSCLC and corresponding normal lung tissues were collected prospectively in the Queen Mary Hospital, University of Hong Kong. Tissue collection protocols were approved by the Joint Hospital and University Institutional Review Board and written informed consents from patients were obtained. Fresh tissues were snap-frozen within 45-60 min after vascular clamping and kept in -70°C until use. Adjacent tumor tissues were fixed in 4% neural buffered formalin for 24 hrs and processed into formalin fixed, paraffin embedded (FFPE) tissue blocks. Tumor classification and differentiation grading was according to the WHO classification of lung tumors, 2004. Tumor typing and pathological staging was performed by a qualified anatomical pathologist (MPW). Clinical parameters and outcomes were charted from hospital records in consultation with relevant clinicians.

### Immunohistochemistry (IHC)

Tissue microarrays were constructed using at least 5 cores of tissue from different representative tumor areas and 1 core of corresponding normal lung from each case. Tumor cores were randomly arranged in the microarray to prevent positional bias during recording of IHC results. 5um thick de-paraffinized tissue microarray sections were subjected to antigen retrieval using microwave heating at 95T°C in 1mM EDTA buffer, pH 8.0. Endogenous peroxidase was quenched with 3% hydrogen peroxide for 10 minutes. Blocked sections were labeled with primary antibodies against NFATc2 (1:50 dilution, Cell Signaling), SOX2 (1:200 dilution, Cell Signaling) and ALDH1A1 (1:1000 dilution, Abcam) overnight at 4°C. Anti-rabbit HRP-labeled polymer (DAKO) was used as a secondary antibody. Color detection was performed by liquid DAB+ substrate chromogen system (DAKO). Primary antibodies were omitted in control reactions. Protein expression levels were semi-quantitatively analyzed using an automated image capturing and analysis system (Aperio).

NFATc2 expression level was scored according to the extent and intensity of nuclear staining in the tumor cells only and expression in the cytoplasm, stromal or inflammatory cells was excluded from evaluation. The intensity was graded as 1, 2, or 3 according to whether nuclear staining was absent or weak, moderate, or strong, respectively. The staining extent was graded as 1, 2, or 3 according to whether expression was observed in scattered individual cells, aggregates of 5 or more but <19 cells, or sheets of 20 or more cells. The products of the 2 grades were then computed, and cases with scores of 4 and above were counted as high level expression.

### Statistics

Data were analyzed by SPSS (version 16.0; SPSS Inc., Chicago, IL, USA), GraphPad Prism 7.0 or Excel (Microsoft, Redmond, WA, USA) software packages and shown as mean ± standard deviations (s.d.). Differential expression between paired tumor/normal tissues were analyzed by Wilcoxon text. Differences between groups were analyzed by *t* test for continuous variables. Differences between growth curve of xenograft model were analyzed by two-way ANOVA. Correlation between NFATc2 and SOX2 mRNA level were analyzed by Pearson correlation test. Correlation between NFATc2, SOX2, ALDH1A1 expressions and clinicopathological variables in lung cancers were analyzed by the χ^2^-test. Association between NFATc2 expression and overall survival and recurrence-free survival were analyzed by the Kaplan–Meier method with log-rank test. Multivariate survival analyses were performed by Cox regression model. Two-sided P values <0.05 were considered as being statistically signi?cant.

## Acknowledgement

We thank the Core Facility and Laboratory Animal Unit of the LKS Faculty of Medicine, The University of Hong Kong for technical support. We are thankful to Dr Terence Kin-Wah LEE, Dr. Stephanie Kwai-Yee Ma and Dr Judy Wai-Ping Yam for their helpful discussion on the study and comments on the manuscript.

## Conflict of interest

We declare there is no conflict of interest amongst any of the listed authors, or between any author and the sponsoring institutions or other parties.

## Figure legends

**Figure 2-figure supplement 1.**
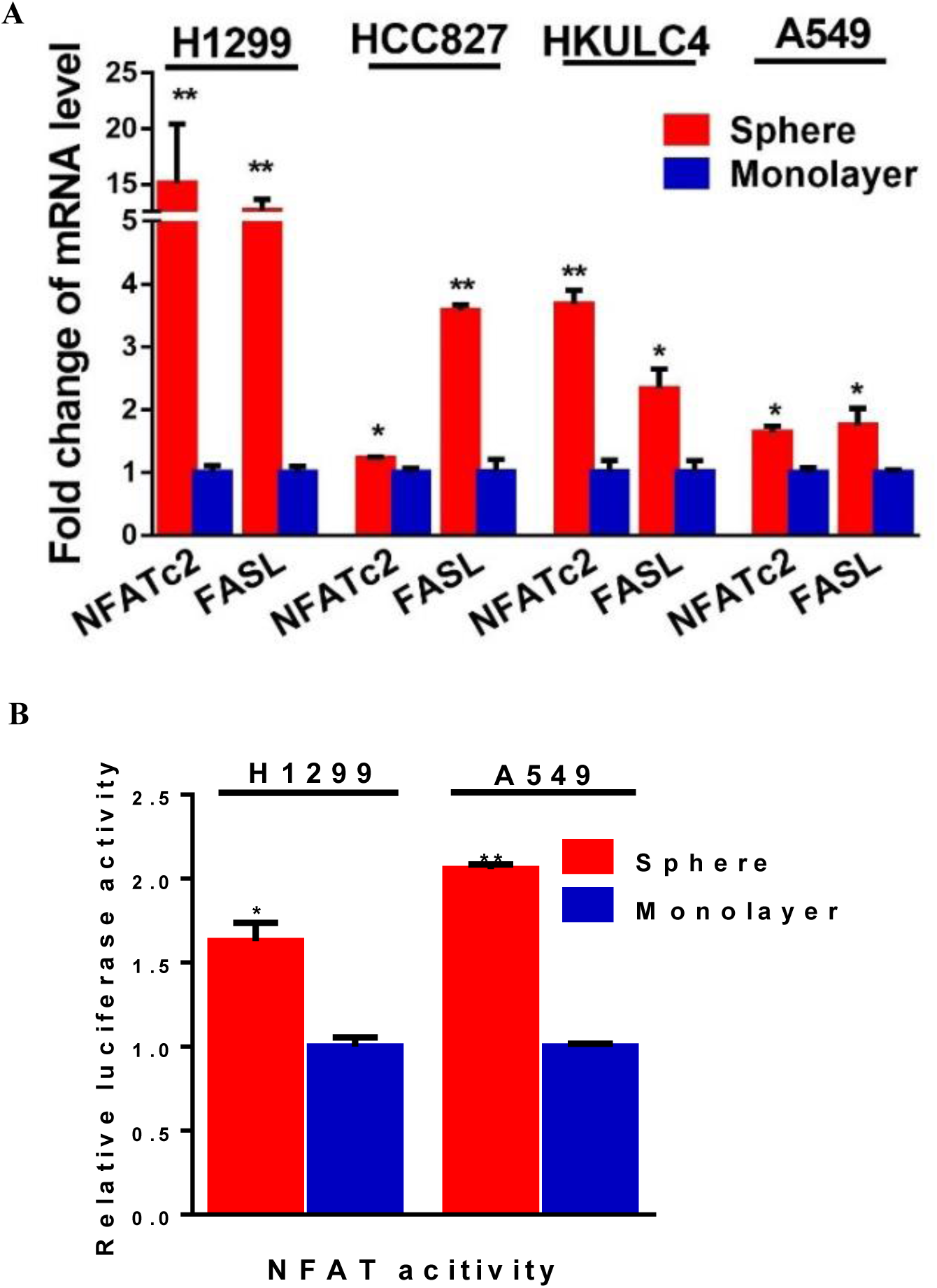
NFATc2 was up-regulated in tumorspheres. **(A)** Expression of *NFATc2* and its target *FASL* analyzed by qPCR in TIC isolated by tumorspheres compared to controls. (**B)** NFAT luciferase reporter activity in TIC isolated by tumorspheres compared to the corresponding monolayer controls. *p<0.05 **p<0.01, comparison with control by t-test. Error bar indicates the mean ± SD for three independent replicates.

**Figure 2-figure supplement 2.**
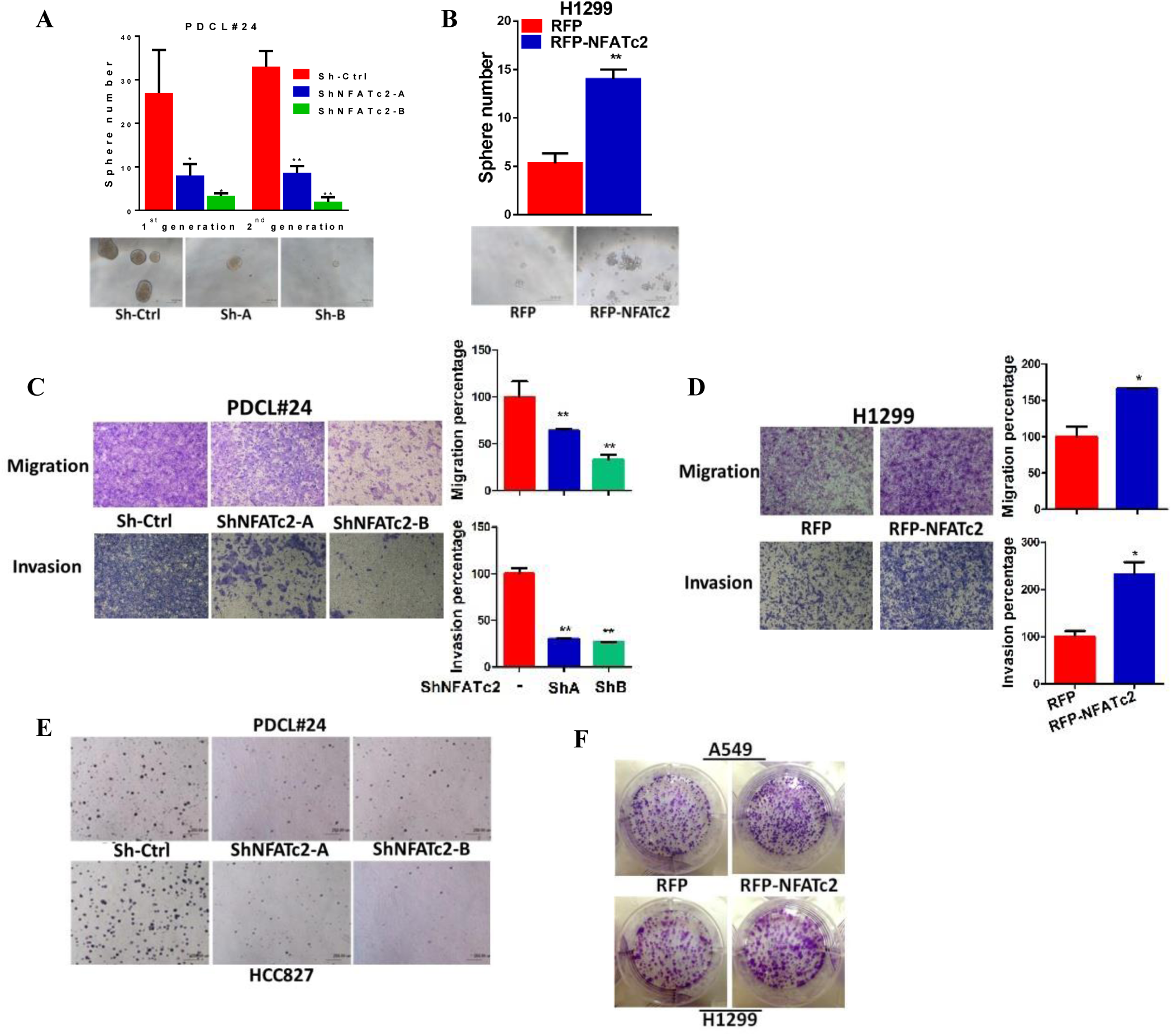
NFATc2 regulated *in vitro* TIC properties. **(A-B)** Tumorsphere formation assays in PDCL#24 cells after stable NFATc2 knockdown (A), or in H1299 cells with stable NFATc2 over-expression (B) compared to controls. **(C-D)** Cell migration and invasion assays in cells with stable NFATc2 knock down (C) or over-expression (D). **(E)** Anchorage independent growth assays of PDCL#24 and HCC827 cells with NFATc2 knockdown. **(F)** Colony formation assay in A549 and H1299 cells with or without NFATc2 overexpression. *p<0.05 **p<0.01, comparison with control by t-test. Error bar indicates the mean ± SD for at least three independent replicates.

**Figure 2-figure supplement 3.**
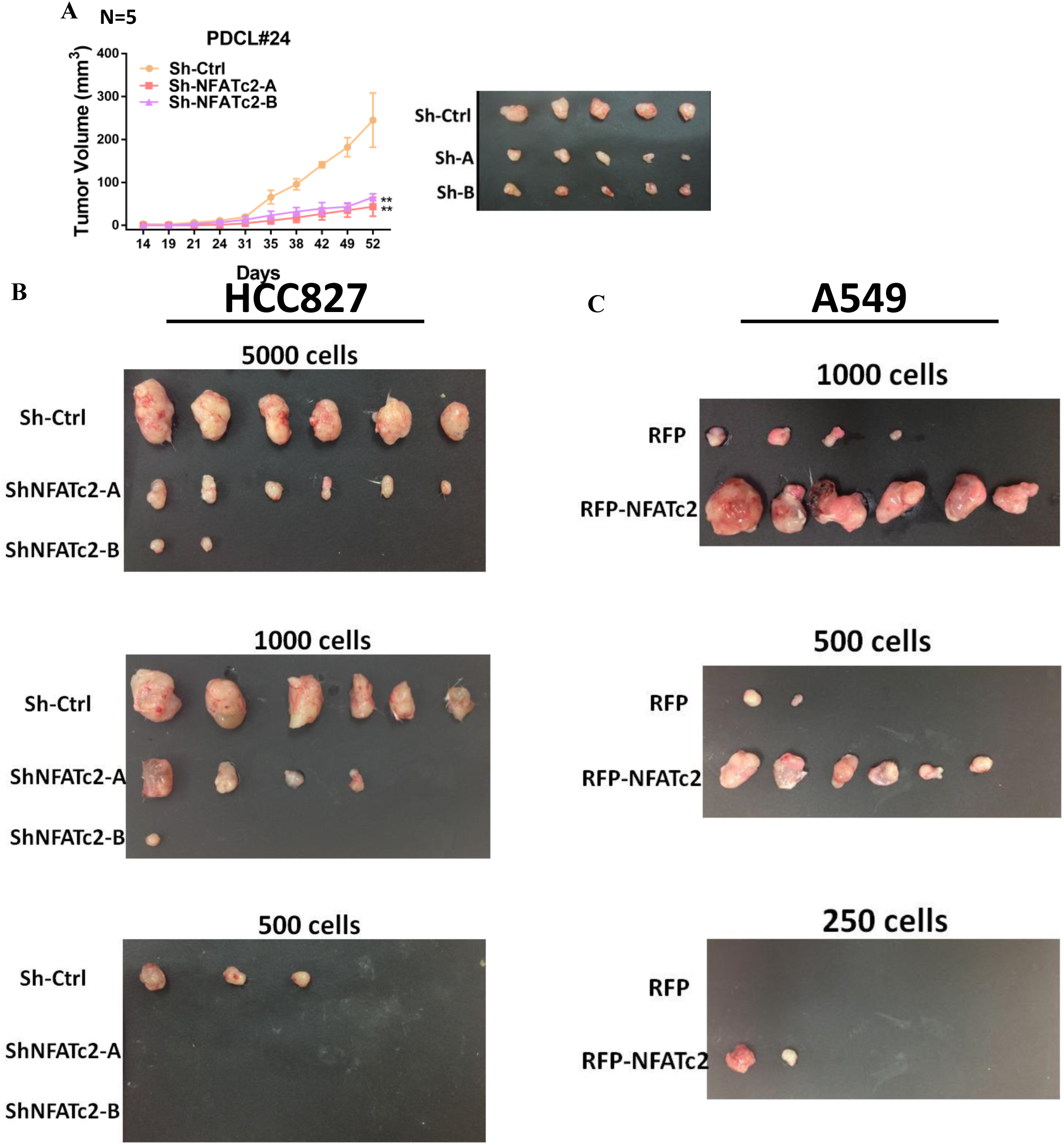
NFATc2 regulated *in vivo* tumorigenesis. **(A)** PDCL#24 cells with or without NFATc2 knockdown were subcutaneously inoculated into the flanks of SCID mice, and tumor volumes were monitored. Representative tumor images and tumor growth curves are shown. Error bars indicate the means ± SD of average tumor volumes of 5 mice. **p<0.0001, comparison with control by two-way ANOVA. Error bar indicates the mean ± SD for average tumor volumes of 5 mice. **(B)** Xenografts of limiting dilution assays of HCC827 cells with stable NFATc2 knockdown. **(C)** Xenografts of limiting dilution assays of A549 cells with NFATc2 over-expression.

**Figure 3-figure supplement 1.**
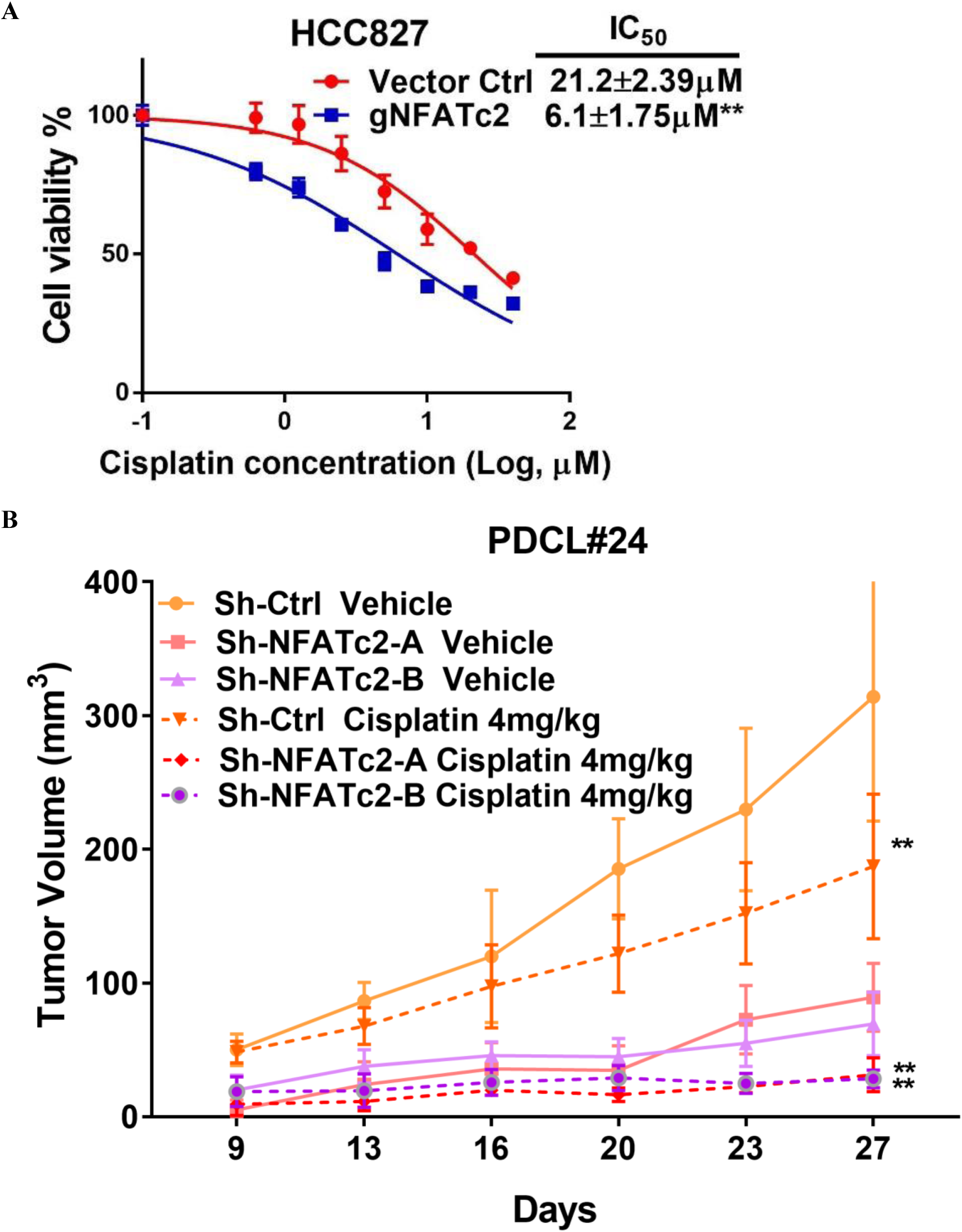
NFATc2 promoted cancer resistance to cisplatin treatment. **(A)** Cisplatin sensitivity analyzed by MTT assay in HCC827 cells with NFATc2 knockout. **p<0.01, comparison with control by t-test. Error bar indicates the mean ± SD for three independent replicates. (**B)** Growth curve of *in vivo* tumor response to cisplatin of PDCL#24 xenografts with or without NFATc2 knockdown. **p<0.0001 versus control vehicle by two-way ANOVA. Error bar indicates the mean ± SD of tumor volumes five mice for Sh-Ctrl vehicle and Sh-NFATc2-A vehicle groups (1 mouse from each group failed to develop tumor) and six mice for other groups.

**Figure 3-figure supplement 2.**
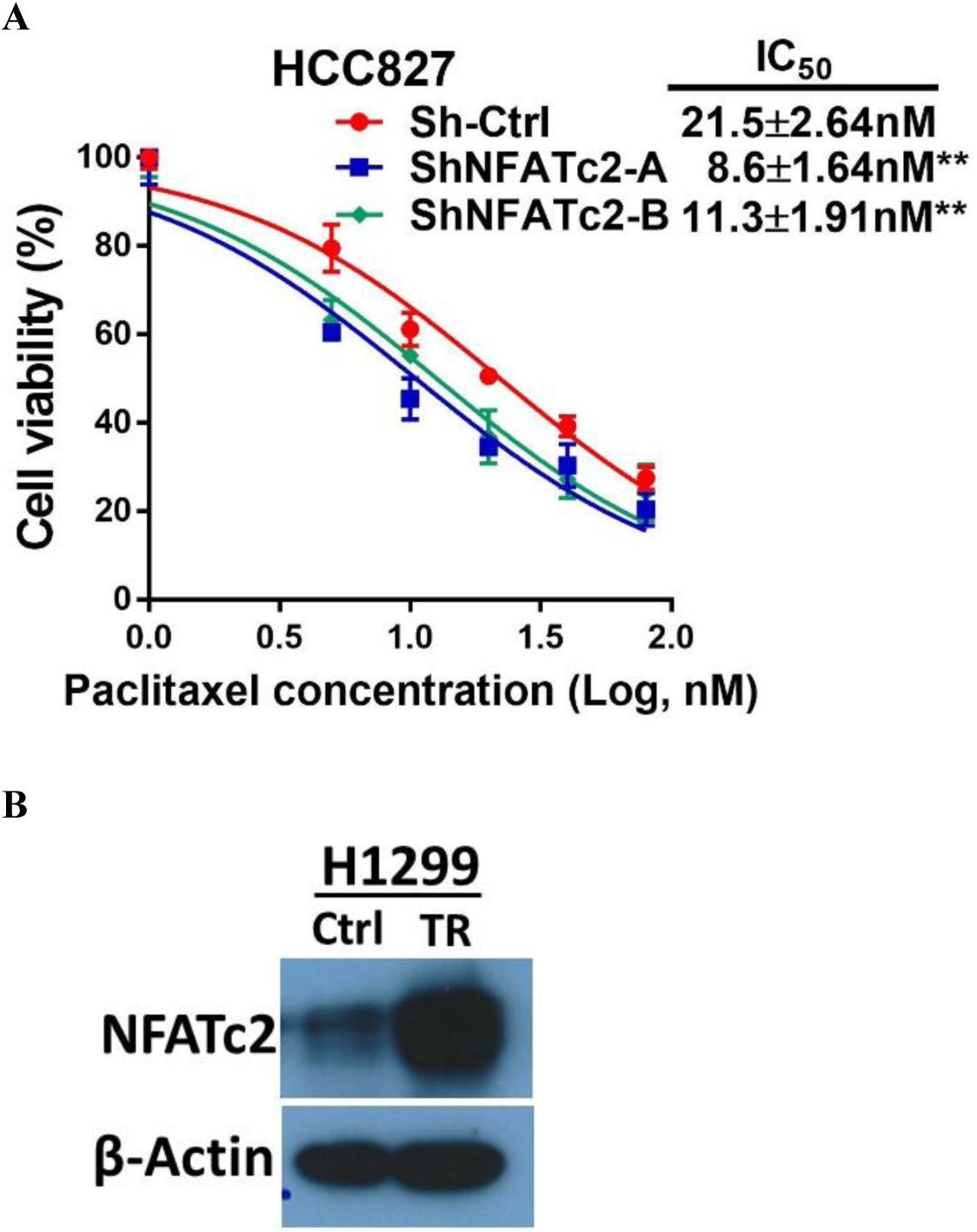
NFATc2 promoted cancer resistance to paclitaxel treatment. **(A)** Paclitaxel sensitivity analyzed by MTT assay in HCC827 cells with NFATc2 knockdown. **p<0.01, comparison with control by t-test. Error bar indicates the mean ± SD for three independent replicates. **(B)** NFATc2 expression analyzed by Western blot in H1299 parental and paclitaxel resistant (TR) cells.

**Figure 3-figure supplement 3.**
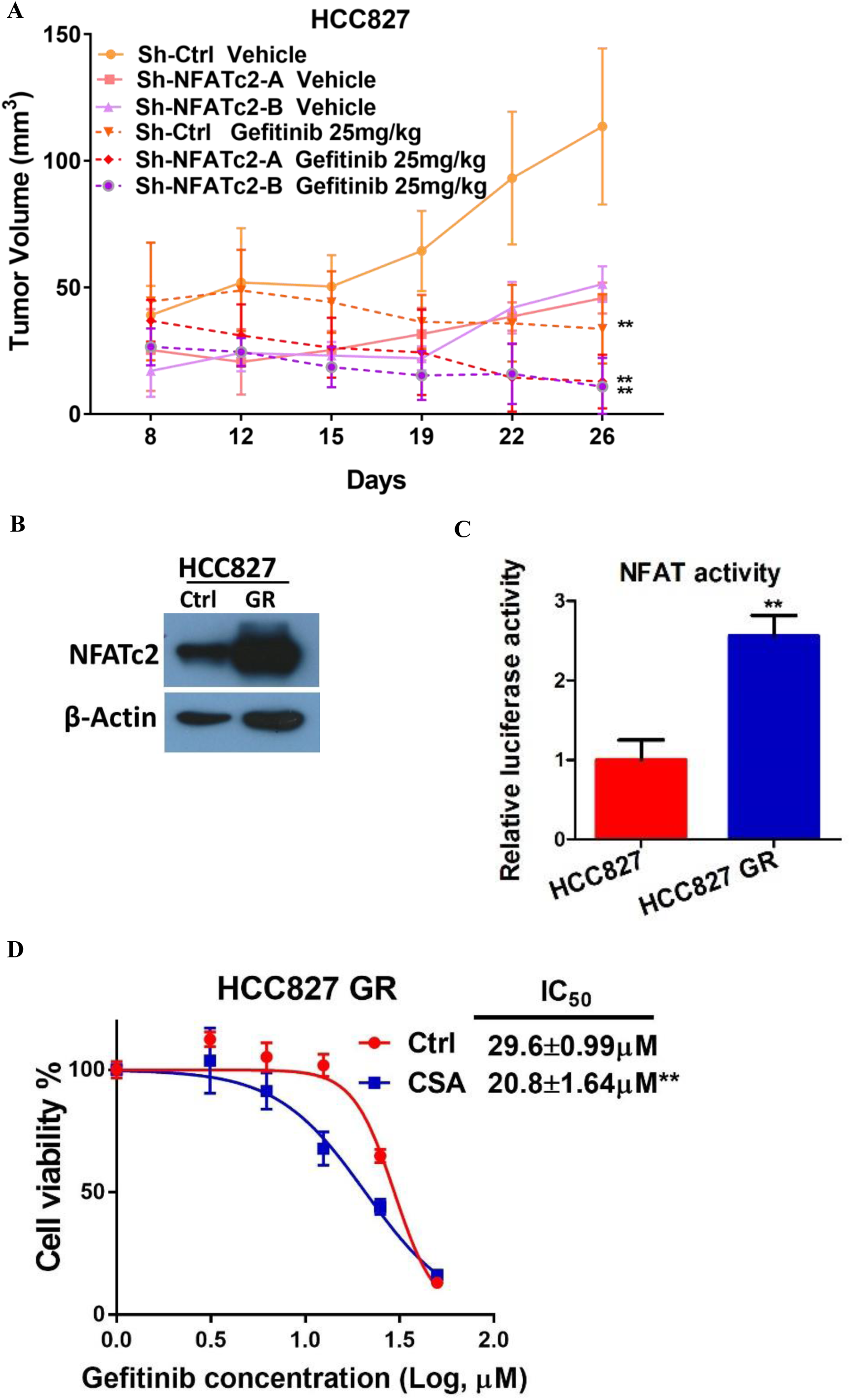
NFATc2 promoted cancer resistance to gefitinib treatment. **(A).** Growth curve of *in vivo* tumor response to gefitinib of HCC827 xenografts with or without NFATc2 knockdown. **p<0.0001 versus control vehicle by two-way ANOVA. Error bar indicates the mean ± SD of tumor volumes of six mice. **(B-C)** NFATc2 expression by Western blot (B) and NFAT activity by luciferase reporter assay (C) in HCC827 parental and gefitinib-resistant (GR) cells. **(D)** Gefitinib sensitivity with or without CSA treatment analyzed by MTT assay in HCC827GR cells. **p<0.01 versus control by Student’s t-test. Error bar indicates mean ± SD for at least three replicates.

**Figure 4-figure supplement 1.**
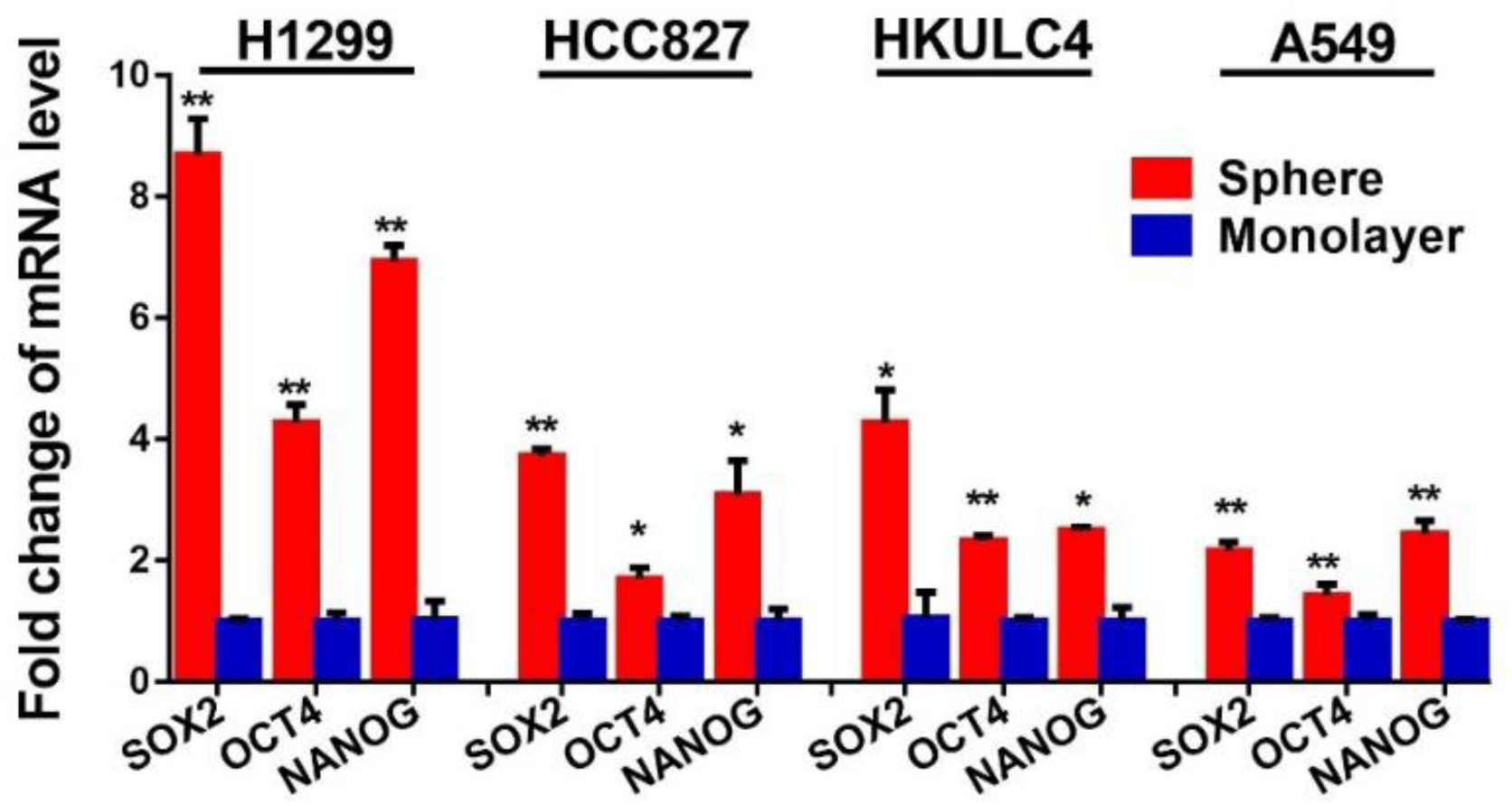
Expression of pluripotency factors in tumorspheres. Expression of pluripotency factors in tumorspheres was analyzed by qPCR and normalized to monolayers of indicated cell lines. *p<0.05, **p<0.01 versus control by t-test. Error bar indicates the mean ± S.D. for three replicates.

**Figure 4-figure supplement 2.**
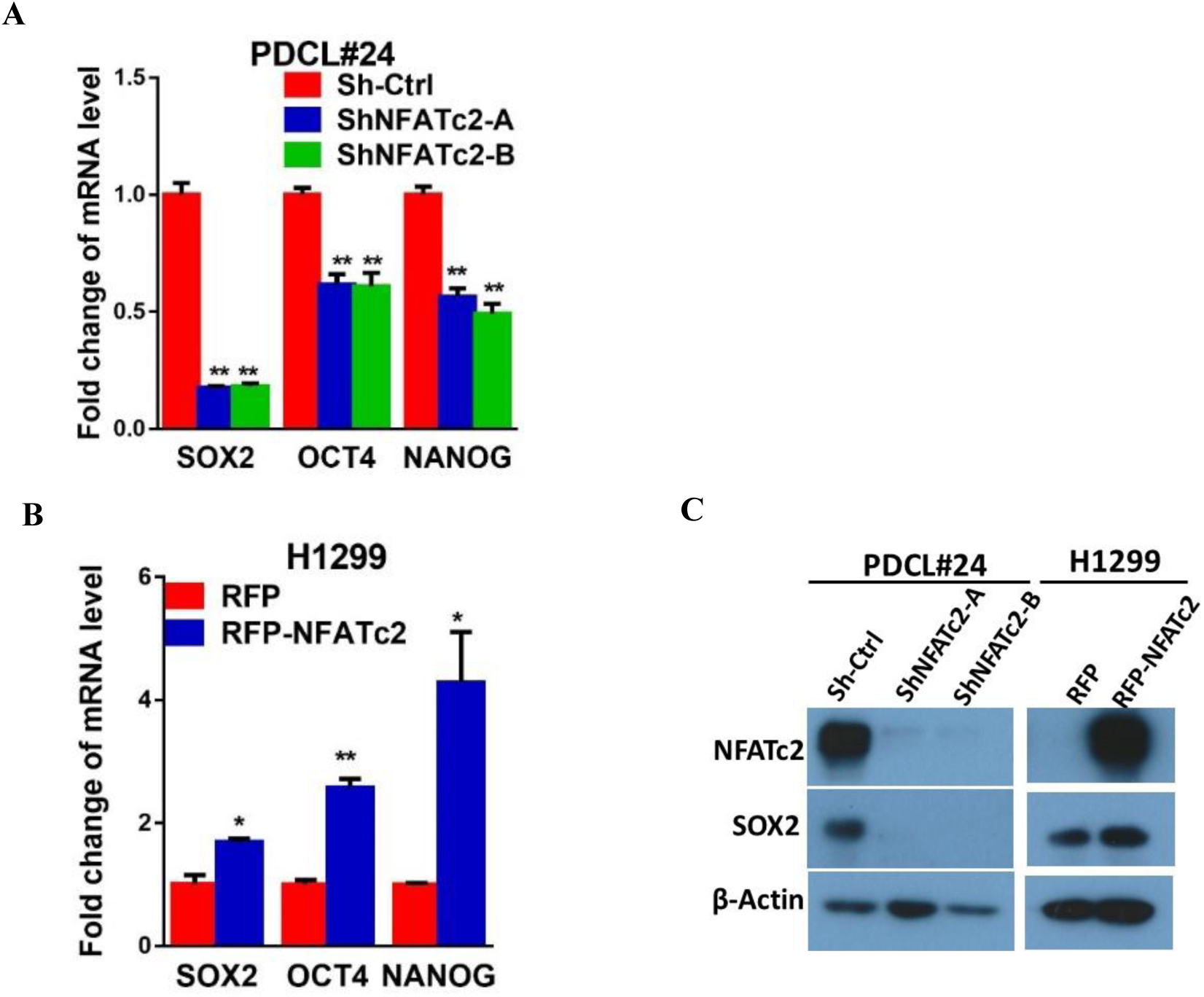
NFATc2 regulated SOX2 expression. **(A-B)** Expressions of *SOX2*, *OCT4*, *NANOG* analyzed by qPCR in PDCL#24 cells with NFATc2 knockdown (A), or H1299 cells with NFATc2 overexpression (B). *p<0.05, **p<0.01 versus control by t-test. Error bar indicates the mean ± S.D. for three replicates**. (C)** Effects of stable NFATc2 knockdown or upregulation on SOX2 expression in lung cancer cells by Western blot analysis.

**Figure 4-figure supplement 3.**
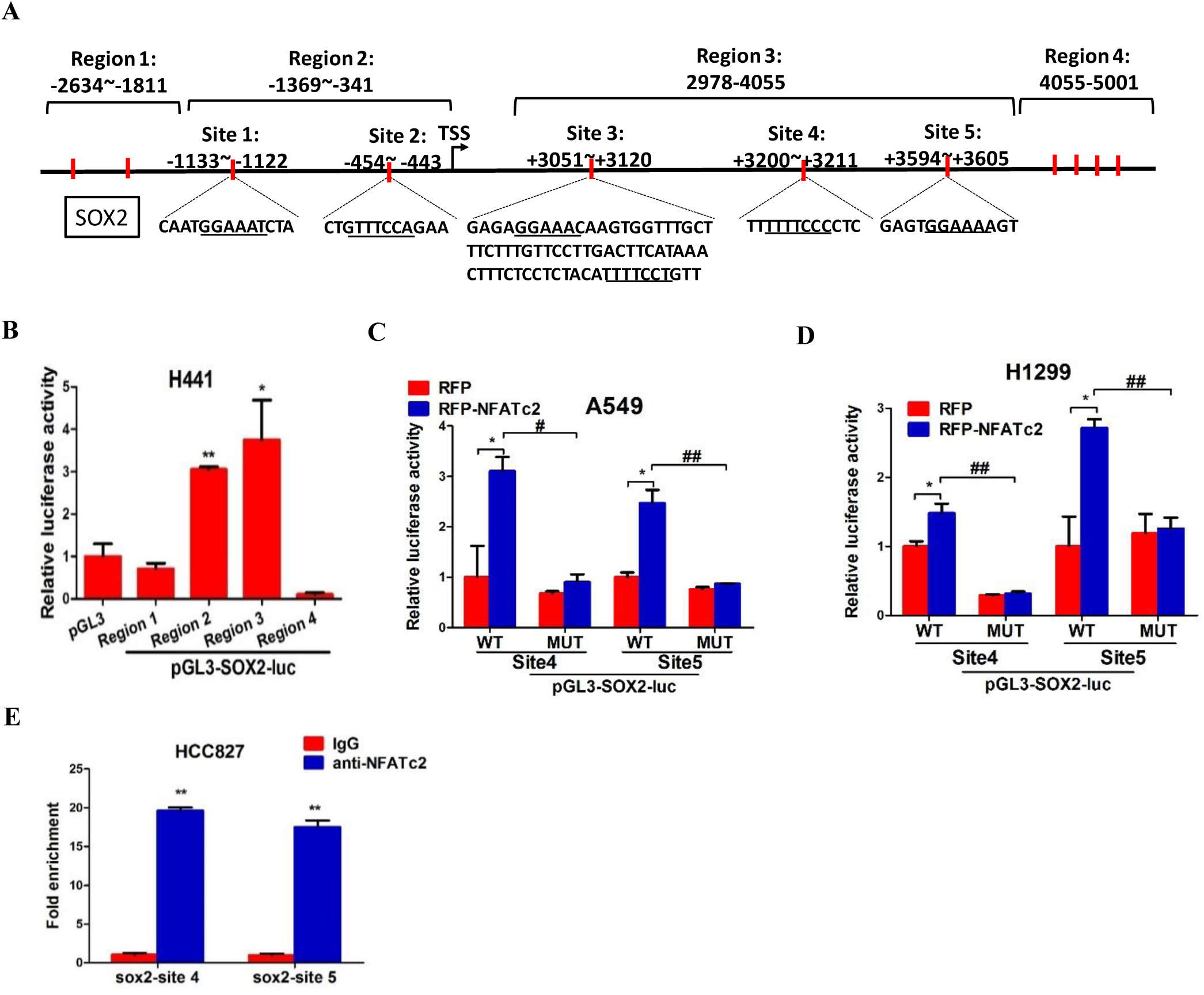
NFATc2 regulated SOX2 expression through binding to 3’ regulatory regions. **(A)** Computational prediction of NFAT binding sites (marked as red curve) on 5’ and 3’ SOX2 regulatory regions (Regions 1 to 4). TSS: transcription start site. **(B)** Transcriptional activity of region 1-4 were studied in H441 cells using luciferase reporter assay. **(C-D)** Luciferase reporter activities of mutant or wild-type SOX2 reporters were analyzed in A549 (C) or H1299 (D) cells with or without NFATc2 stable overexpression. *p<0.05, **p<0.01, comparison with RFP; **^#^** p<0.05, ^##^ p<0.01, wild type versus mutant in RFP-NFATc2 cells by t-test. Error bars indicate the mean ± SD for at least three independent replicates. **(E)** Confirmation of NFATc2 binding to candidate SOX2 sites by ChIP-qPCR analysis in HCC827 cells. For B and E, *p<0.05, **p<0.01 versus control by Student’s t-test. Error bars indicate the mean ± SD for at least three independent replicates.

**Figure 4-figure supplement 4.**
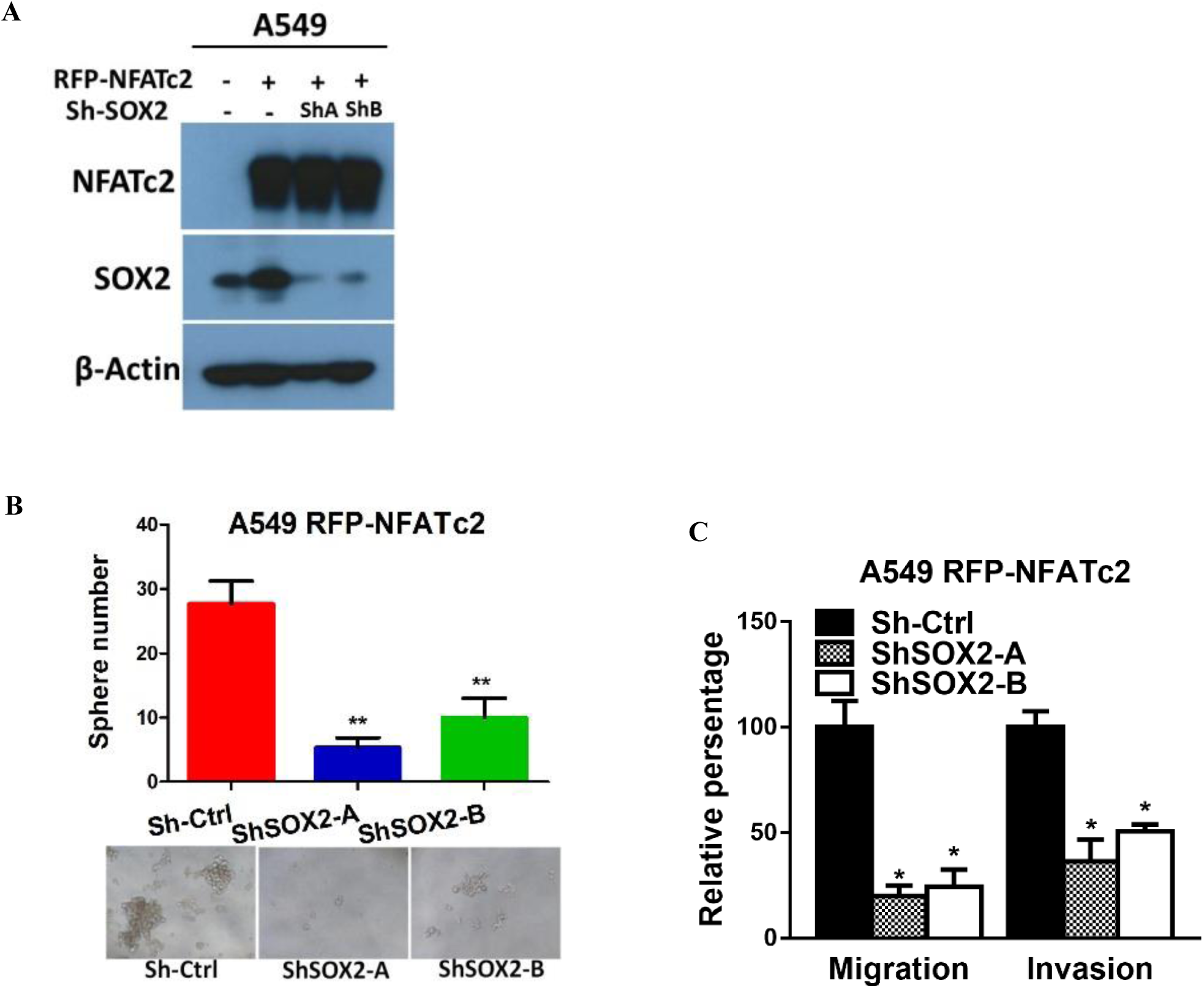
NFATc2 regulated tumor function through SOX2. **(A)** Expression of NFATc2 and SOX2 analyzed by Western blot in A549 cells with or without NFATc2 overexpression and SOX2 stable knockdown. **(B-C)** Effect of SOX2 knockdown on tumorsphere formation (B) and cell migration and invasion ability (C) of NFATc2 overexpressing A549 cells. *p<0.05, **p<0.01 versus control by t-test. Error bars indicate the mean ± SD for at least three independent replicates.

**Figure 5-figure supplement 1.**
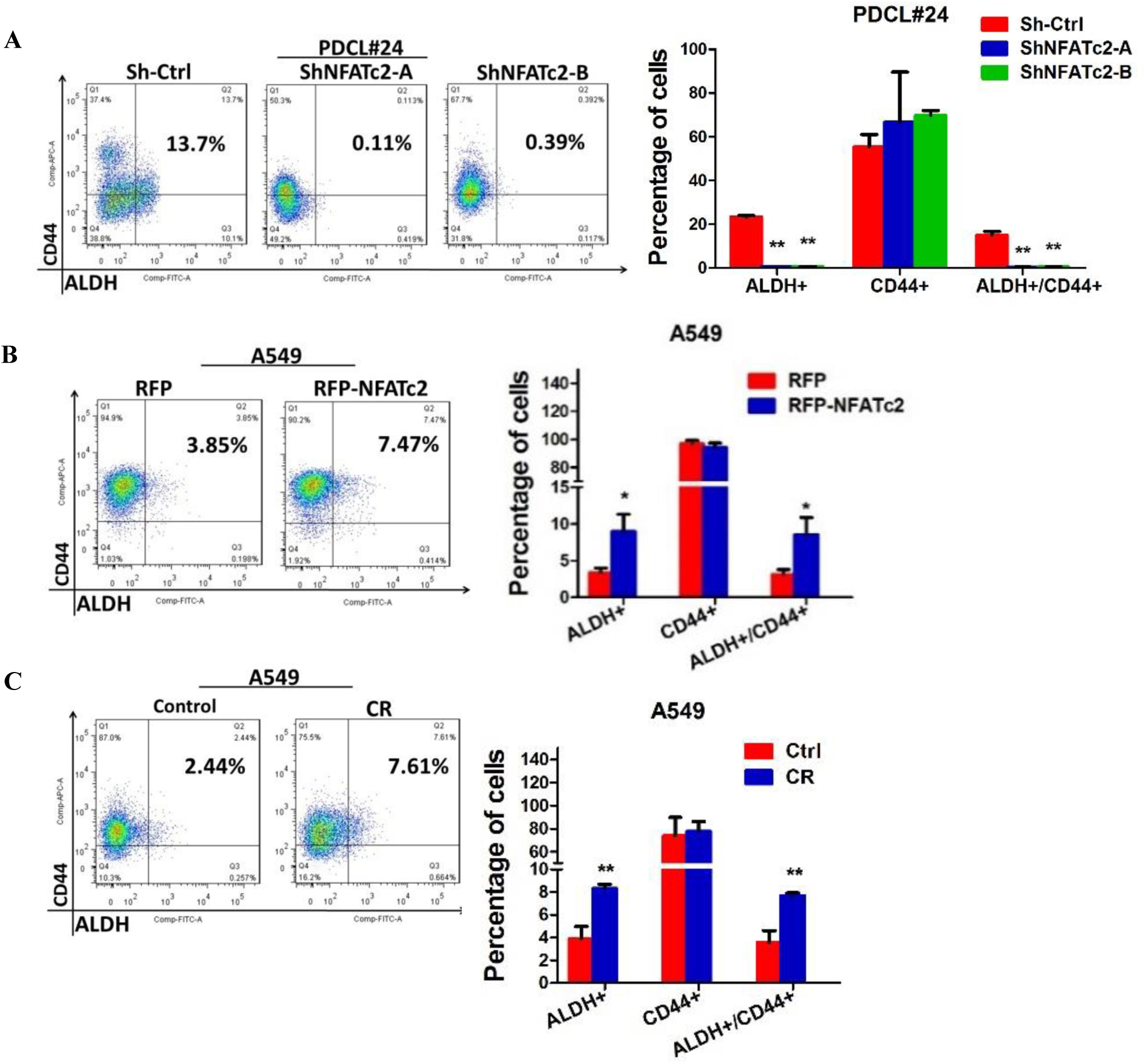
NFATc2 regulated the ALDH^+^ population. **A-C.** Flow cytometry analysis of ALDH/CD44 distribution in PDCL#24 cells with stable NFATc2 knockdown (A), A549 cells with NFATc2 overexpression (B), or A549CR compared to parental cells (C). *p<0.05, **p<0.01 versus control by t-test. Error bars indicate the mean ± SD for at least three independent replicates.

**Figure 5-figure supplement 2.**
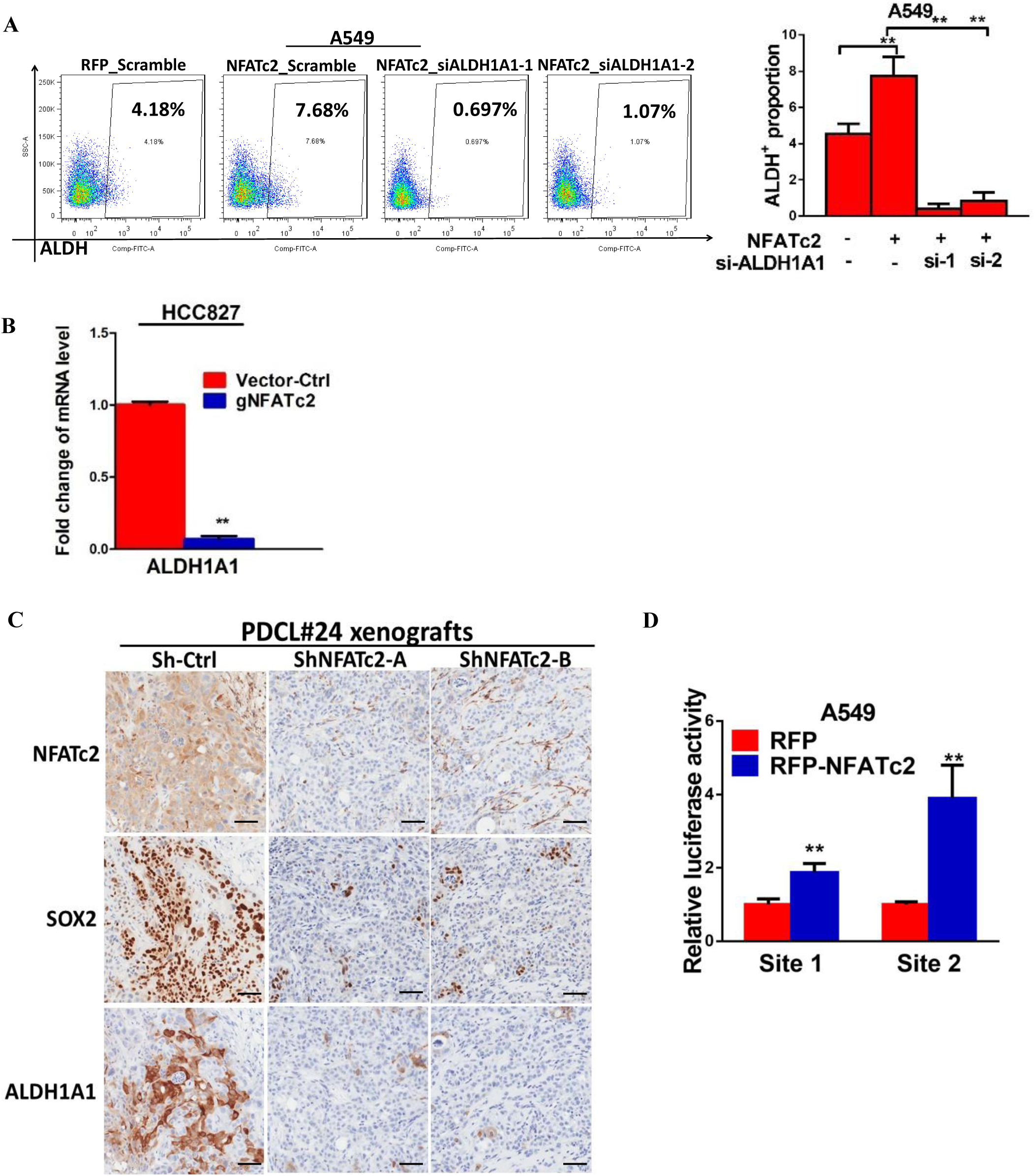
NFATc2 regulated ALDH1A1 expression. **(A)** ALDH^+^ proportions analyzed by flow cytometry in A549 cells with NFATc2 overexpression and transient ALDH1A1 knockdown. **(B)** mRNA level of *ALDH1A1* analyzed by qPCR in HCC827 cells with NFATc2 knockout. **(C)** Representative images of immunohistochemical expression of NFATc2, SOX2 and ALDH1A1 in xenografts derived from PDCL#24 cells with or without NFATc2 knockdown. Scale bars, 50 µM. **(D)** Luciferase reporter assay for site1 and 2 in A549 cells with or without NFATc2 overexpression. **p<0.01 versus control by t-test. Error bars indicate the mean ± SD for at least three independent replicates.

**Figure 5-figure supplement 3.**
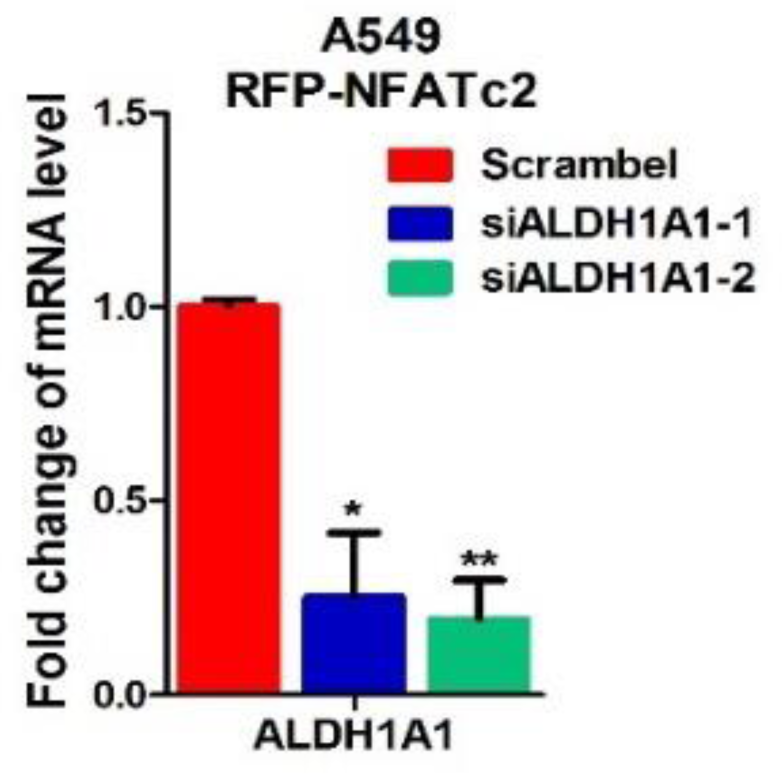
Effect of siALDH1A1 on ALDH1A1 expression. Analysis of *ALDH1A1* expression by qPCR in NFATc2-overexpressing A549 cells with ALDH1A1 knockdown. **p<0.01 versus control by t-test. Error bar indicates the mean ± S.D. for 3 replicates.

**Figure 5-figure supplement 4.**
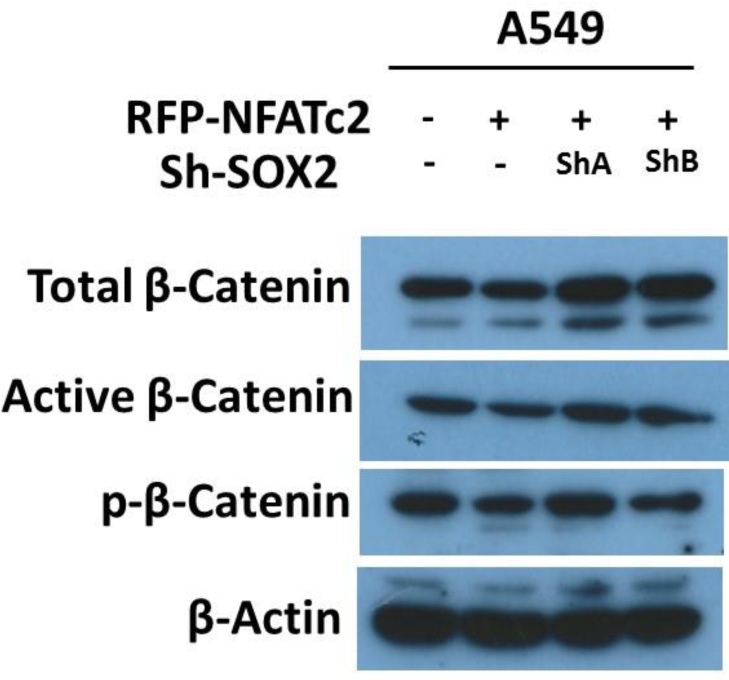
Effect of NFATc2/SOX2 on β-catenin activity. Expressions of total β-catenin, active β-catenin (non-phosphorylated), and phosphorylated β-catenin (p-β-catenin) analyzed by immunoblot in A549 with or without NFATc2 overexpression and SOX2 knockdown.

